# Characterization of prefusion-F-specific antibodies elicited by natural infection with human metapneumovirus

**DOI:** 10.1101/2022.03.28.486060

**Authors:** Scott A. Rush, Gurpreet Brar, Ching-Lin Hsieh, Emilie Chautard, Jennifer N. Rainho-Tomko, Chris Slade, Christine A. Bricault, Ana Kume, James Kearns, Rachel Groppo, Sophia Mundle, Linong Zhang, Danilo Casimiro, Tong-Ming Fu, Joshua M. DiNapoli, Jason S. McLellan

## Abstract

Human metapneumovirus (hMPV) is a major cause of acute respiratory tract infections in infants and the elderly for which there are no approved vaccines or antibody therapies. The viral fusion (F) glycoprotein is required for entry and is the primary target of neutralizing antibodies, however, little is known about the humoral immune response generated by humans as a result of natural infection. Here, we use stabilized hMPV F proteins to interrogate memory B cells from two elderly donors. We obtained over 700 paired non-IgM antibody sequences representing 563 clonotypes, indicative of a highly polyclonal antibody response to hMPV F in these individuals. Characterization of 136 of these monoclonal antibodies revealed broad recognition of the hMPV F surface, with potent neutralizing antibodies targeting each antigenic site. Cryo-EM structures of two neutralizing antibodies reveal the molecular basis for recognition of two prefusion-specific epitopes at the membrane-distal apex of hMPV F. Collectively these results provide new insights into the humoral response to hMPV infection in the elderly and will guide development of novel vaccine antigens.

## INTRODUCTION

Human metapneumovirus (hMPV) is an enveloped negative-sense RNA virus of the *Pneumoviridae* family that has been circulating for at least 70 years but was only identified in 2001 (van den Hoogen et al., 2001). Analysis of hMPV genetic lineages identifies two genotypes, A and B (Peret et al., 2002; van den Hoogen et al., 2004). The genotypes are further divided into 5 subgroups (A1, A2a, A2b, B1 and B2) primarily based upon the attachment protein sequence variability (Nao et al., 2020; van den Hoogen et al., 2004; Yang et al., 2013). hMPV has a seasonality of winter and early spring with co-circulation of the two genotypes frequently observed (Kahn, 2006). It is now recognized as a leading cause of respiratory tract infections in infants with near universal pathogen exposure by the age of five (van den Hoogen et al., 2001; Wang et al., 2021). In 2018, it was estimated that the global burden of hMPV-associated acute lower respiratory tract infections in children under five was 14 million cases with approximately 643,000 associated hospitalizations (Wang et al., 2021). Despite early exposure, reinfection continues throughout life with symptoms that are generally mild. However, reinfections among the immunocompromised and the elderly can lead to more severe disease such as bronchiolitis and pneumonia (Boivin et al., 2002; Boivin et al., 2007; Walsh et al., 2008). Although vaccines and monoclonal antibodies are in various stages of early clinical development (Biacchesi et al., 2005; Chupin et al., 2021; Cox et al., 2014; Cseke et al., 2007; Hamelin et al., 2007; Herfst et al., 2008; Huang et al., 2021; Karron et al., 2018; Levy et al., 2013; Olmedillas et al., 2018; Schuster et al., 2015; Stepanova et al., 2020; Stewart-Jones et al., 2021), there are currently no FDA-approved interventions available. To develop an effective hMPV countermeasure, it will be important to have a thorough understanding of the humoral immune response that is elicited by natural infection.

Like other pneumoviruses, such as respiratory syncytial virus (RSV), the envelope of the virion is decorated with two glycoproteins: the fusion (F) glycoprotein and the attachment (G) glycoprotein. hMPV F and G facilitate viral attachment through interactions with cellular glycosaminoglycans (Chang et al., 2012; Huang et al., 2021; Klimyte et al., 2016; Thammawat et al., 2008). Additionally, F may interact with RGD-binding integrins through a highly conserved RGD motif (Cox et al., 2012; Cseke et al., 2009; Wei et al., 2014). However, whereas G is dispensable for propagation in cell culture (Biacchesi et al., 2004; Dubois et al., 2019), F is required for viral entry because it facilitates fusion of the viral and host-cell membranes (Mas and Melero, 2013; Melero and Mas, 2015). Although the immune response elicited by hMPV G has been found to be nonprotective, hMPV F elicits a potent neutralizing antibody response (Biacchesi et al., 2004) and is therefore a major focus of vaccine development efforts.

Like all class I viral fusion glycoproteins, hMPV F is initially translated as an inactive precursor (F_0_). Activation requires proteolytic cleavage by a host-cell protease at a conserved RQSR sequence located between the N-terminal F_2_ subunit and the C-terminal F_1_ subunit. This cleavage is likely performed by serine proteases such as TMPRSS2 (Shirogane et al., 2008) on the target cell, although some F protein may be cleaved during viral egress. Following proteolysis, the disulfide-linked F_2_/F_1_ heterodimers trimerize to adopt the metastable prefusion (preF) conformation (Battles et al., 2017). In response to an unknown stimulus or thermodynamic instability, the preF protein undergoes a dramatic rearrangement of its tertiary structure, extending and inserting the hydrophobic fusion peptide at the N-terminus of the F_1_ subunit into the host-cell membrane. This elongated pre-hairpin intermediate then collapses back upon itself to bring the F_1_ N- and C-termini, and thus the host-cell and viral membrane, into close proximity, resulting in the fusion of the two membranes and adoption of the highly stable postfusion (postF) conformation (Mas et al., 2016).

Through large-scale antibody isolation and characterization studies of RSV F, six major antigenic sites (Ø, I, II, III, IV, and V) covering the preF protein surface have been described (Gilman et al., 2016). Antigenic sites Ø and V are preF-specific as they comprise secondary structure elements that undergo significant rearrangement during the preF-to-postF transition (Gilman et al., 2016; McLellan et al., 2013). Antibodies that recognize site III generally have a higher affinity to preF than postF and are thus considered preF-preferring (Goodwin et al., 2018). Sites II and IV are found on both preF and postF (McLellan, 2015; Swanson et al., 2011), whereas site I antibodies have a higher affinity for postF (Gilman et al., 2016). RSV and hMPV F proteins share ∼33% sequence identity (van den Hoogen et al., 2002), and structures of the two proteins in their prefusion conformations are similar (Battles et al., 2017; McLellan et al., 2013). Indeed, several previously isolated RSV F-reactive antibodies are cross-reactive with hMPV F, including MPE8, 101F, and M1C7, which recognize sites III, IV, and V, respectively (Corti et al., 2013; Wu et al., 2007; Xiao et al., 2019). Additionally, hMPV F-specific antibodies DS7 and 338 recognize sites I and II (Ulbrandt et al., 2008; Ulbrandt et al., 2006; Wen et al., 2012; Williams et al., 2007), and a human antibody, MPV458, binds hMPV F site Ø within the trimer interface at an epitope that is inaccessible on closed preF trimers (Huang et al., 2020). Although these hMPV F-directed antibodies have helped provide some insight into the antigenicity of hMPV F, there remains a need for a comprehensive understanding of the hMPV F-directed antibodies elicited by natural infection.

Here, we describe a high-throughput antibody isolation effort using memory B cells from two elderly, naturally hMPV infected donors. From these cells >100 monoclonal antibodies were produced and assessed for reactivity to preF and postF antigens, neutralizing activity, and antigenic site recognition. Furthermore, single-particle cryo-EM structures were obtained for two neutralizing antibodies targeting the preF-specific antigenic sites Ø and V, providing the first structural information on hMPV F antibodies targeting these sites. Collectively, the antibodies isolated and characterized here will help guide development of hMPV F vaccines and immunotherapeutics.

## RESULTS

### Screening and prioritization of donors for antibody discovery

Sanofi Pasteur VaxDesign maintains a cohort of PBMC and plasma from human donors for use in pre-clinical research studies. We screened a total of 31 plasma samples from donors between the ages of 59 and 76 under the assumption that these individuals had long histories of repeated exposure to hMPV and were likely to have a robust memory B cell pool for antibody discovery. Donor and donation details can be found in **Table S1**. Donor plasma were initially tested in an ELISA against hMPV preF A1 strain NL/1/00 (Battles et al., 2017; Stewart-Jones et al., 2021), with titers ranging from 4,000 to 64,000 (**Table S1,** **Figure 1A**). We expanded analysis of Donor 2.3 and 4.2 to include ELISA titers against hMPV A1 postF (**Fig 1B**) and determination of virus microneutralization titer against hMPV A2 CAN97-83 (**Fig 1C**). We also performed flow cytometry analysis on cryopreserved PBMCs from these donors (**Fig 1D**), through which we observed an appreciable frequency of hMPV A1 preF-specific memory B cells. Specifically, we observed that 0.50% and 0.42% of class-switched B cells (CD3^-^, CD8^-^, CD14^-^, CD19^+^, CD20^+^, IgM^-^) bound the preF probe in Donor 2.3 and Donor 4.2, respectively. Based on these observations, we decided to proceed with single B cell sequencing from these two donors.

**Figure 1.**
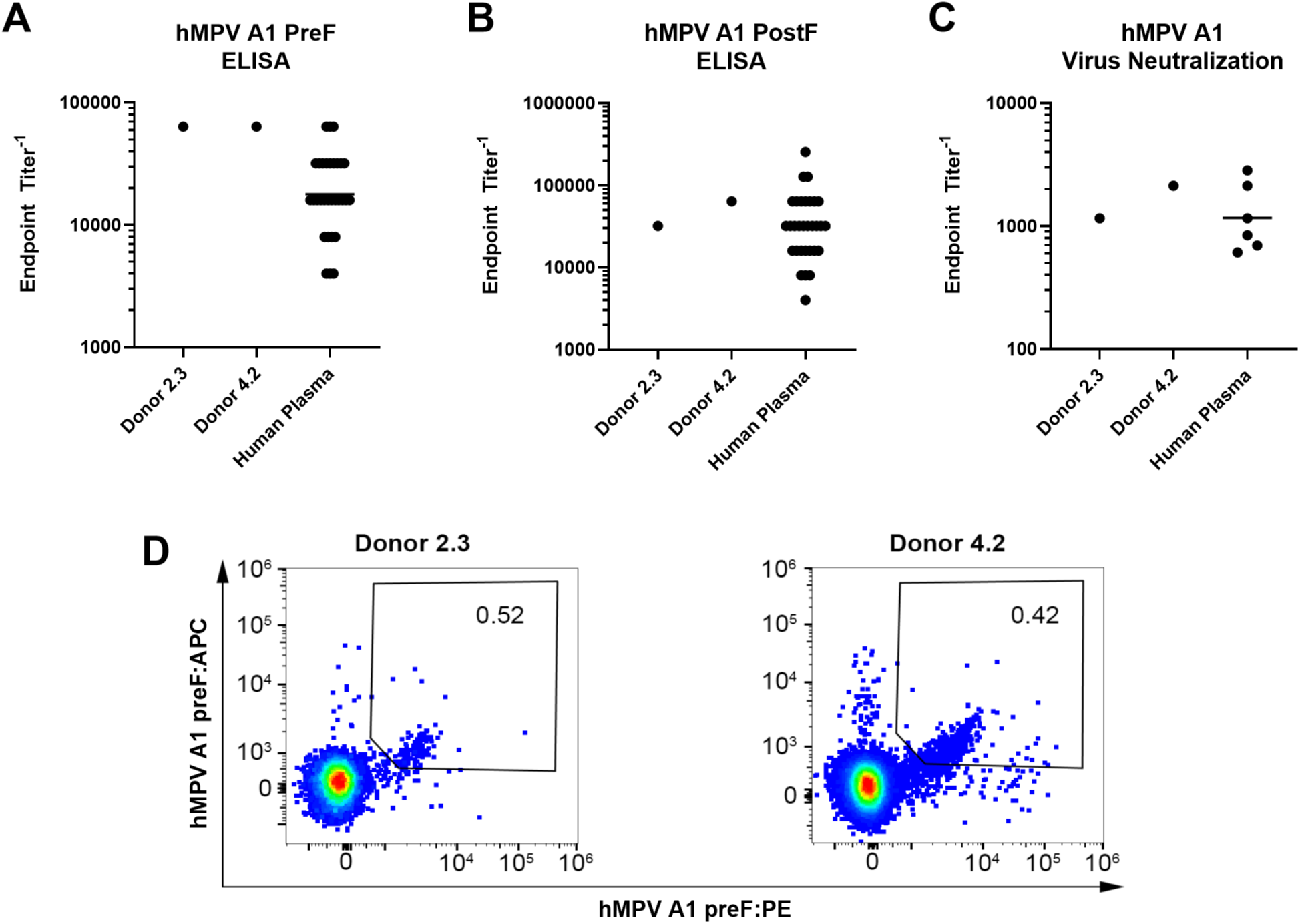
Characterization of sera and PBMCs from hMPV-immune elderly donors used for antibody discovery. **A.** hMPV A1 prefusion F serum ELISA titer; **B.** hMPV A1 postfusion F serum ELISA titer; **C.** hMPV B2 prefusion F serum ELISA titer. For each serological assay, all human plasma samples tested in that assay are shown in the third column; **D.** Frequency of sorted hMPV preF-specific IgM-B cells in Donor 2.3 and Donor 4.2.

### Identification of hMPV preF-reactive antibodies by memory B cell sequencing and analytics

We proceeded with sequencing of memory B cells from Donors 2.3 and 4.2. We sorted a total of 8,573 preF-binding B cells from Donor 2.3 and 4,326 pre-F-binding B cells from Donor 4.2 using a fluorescently tagged hMPV preF protein (115-BV-DS) as a probe. The mRNA from individual B cells was bar coded and sequenced, with resulting paired heavy and light chain sequences annotated and analyzed as described in the Methods. We obtained a total of 359 and 381 paired IgG, IgD and IgA sequences from Donors 2.3 and 4.2, respectively (**Fig 2A**). The sequences from Donor 2.3 could be separated into 235 lineages representing 290 different clonotypes. The sequences from Donor 4.2 could be separated into 231 lineages representing 273 different clonotypes.

**Figure 2.**
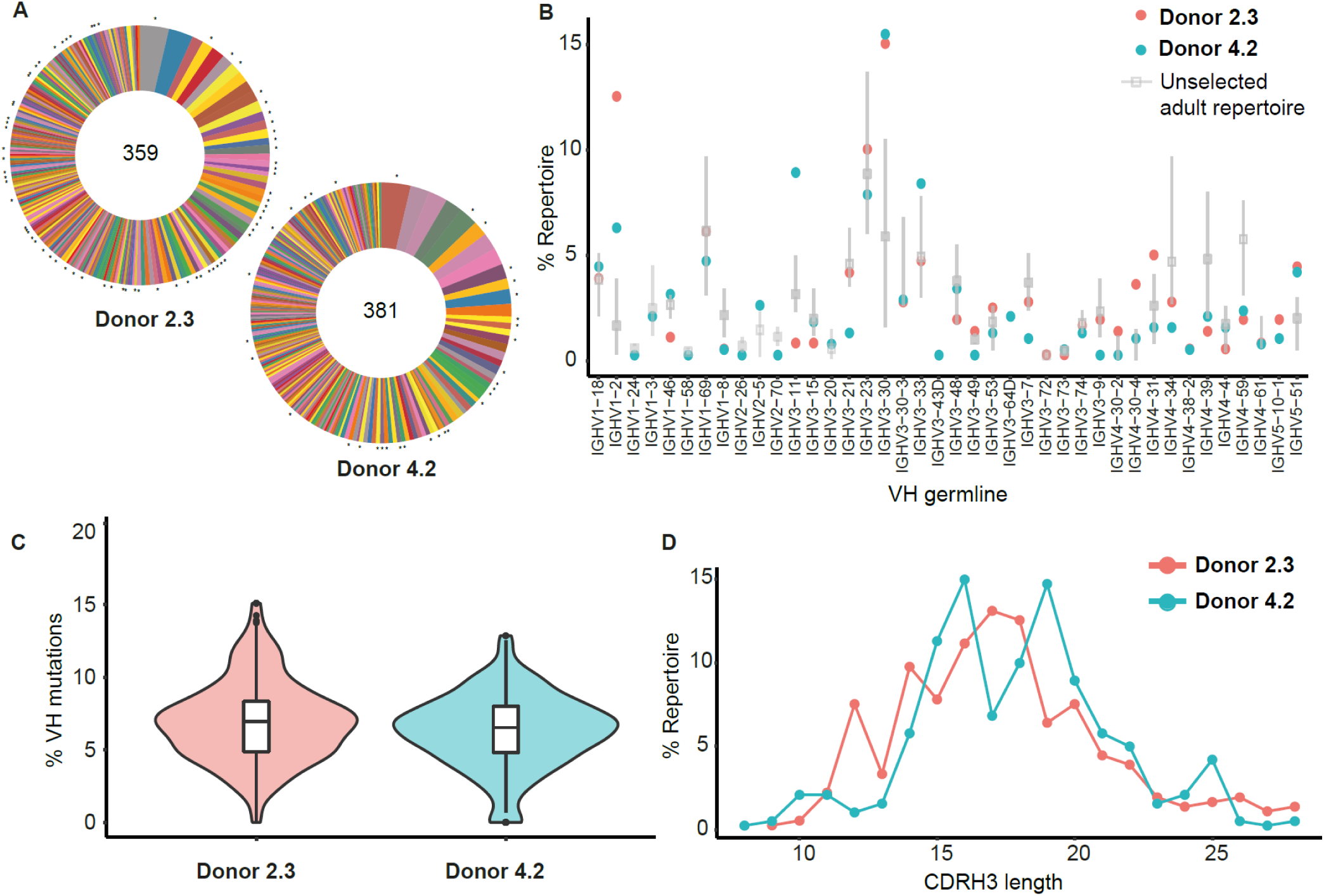
Repertoire analysis of Donors 2.3 and 4.2. Paired heavy and light chain antibody data for Donors 2.3 and 4.2 were individually analyzed using the Immcantation pipeline as described in the methods. Only paired non-IgM sequences are displayed. **A.** Heavy chain lineages for each donor. The size of each slice is proportional to the size of the lineage, and the number in the center represents the number of sequences analyzed for each donor. Asterisks indicate lineages for which at least one mAb was expressed and characterized. **B.** VH germline frequency in donors, with comparison to a previously published adult repertoire derived from 11 healthy individuals. **C.** Violin plot indicating the nucleotide substitution frequency in the VH region of mAbs from each donor, excluding CDRH3 and framework 4. The width is proportional to the fraction of antibodies with the indicated mutation frequency. **D.** CDRH3 length frequency in each donor.

We next explored IGHV usage in these two donors compared to repertoires of previously published human donors (**Fig. 2B**) (Boyd et al., 2010). We found that both donors used IGHV 1-2, 3-30, and 5-51 at a higher frequency than the previously published repertoires. Additionally, Donor 2.3 used IGHV 4-30-4 and 4-31 while Donor 4.2 used IGHV 2-5, 3-11, and 3-33 at a higher frequency than previously published donors. Both donors had similar VH nucleotide substitution frequencies with Donor 2.3 having a median of 6.9% and range 0–15.1% and Donor 4.2 having a median of 6.5% and range 0–12.9% (**Fig. 2C**). Finally, we assessed CDRH3 length in both donors and found that they had a similar distribution, with Donor 2.3 having a majority of 17 to 18 amino-acid-long CDRH3s and Donor 4.2 having a majority of 16 and 19 amino-acid-long CDRH3s (**Fig. 2D, Table S2**).

### Expression and characterization of monoclonal antibodies

We selected 100 paired heavy and light chain sequences to express from Donor 2.3, and 36 sequences to express from Donor 4.2. For Donor 2.3, the 100 antibodies expressed represented 81 clonal groups and 72 distinct heavy chain lineages, with 5 of the top 10 lineages having at least 1 representative. For Donor 4.2, the 36 sequences expressed represented 34 clonal groups and the same number of distinct heavy chain lineages, with 5 of the top 10 lineages having at least 1 representative (**Fig. 2A, Table S2**). Sequences were selected to obtain diversity in lineage coverage, V gene usage, and CDRH3 length. In addition, some sequences were selected based on CDRH3 features deemed to be of particular interest (*e.g.*, cysteine content). All antibodies were expressed as human IgG1 and characterized by ELISA against preF derived from hMPV subgroups A1(NL/1/00) and B2(TN99-419). In addition, all antibodies were tested for their capacity to neutralize hMPV A2 strain CAN97-83 (**Table S3**). Of the 136 antibodies expressed, 115 demonstrated binding to hMPV F by either ELISA or Octet (84.5% hit rate), and 106 demonstrated binding by ELISA only (titers ≥333 ng/ml were considered non-binding). We sometimes observed disparity between antigen binding by ELISA and Octet, presumably due to differences in antigen presentation on the surfaces. Despite performing the B cell sort with hMPV A1 preF, the vast majority of F-specific antibodies obtained were cross-reactive with A1 and B2 preF (**Table S3**). Only 1 antibody (SAN27-60) appeared to be A1 specific based on ELISA. Interestingly, 32 of 136 antibodies appeared to demonstrate a binding preference for B2 preF based on ELISA binding >100-fold. Of the 115 preF-binding antibodies expressed across both donors, 94 had detectable neutralization (81.7%) at ≤12.5 µg/mL and 14 (12.2%) demonstrated a potent neutralization titer (FRNT_50_, mAb concentration resulting in 50% reduction in number of foci) of ≤0.1 µg/mL that compared favorably to published antibodies DS7, 338, MPE8, and 101F (**Table S3**) (Corti et al., 2013; Mas et al., 2016; Ulbrandt et al., 2006; Wen et al., 2012). We observed a similar distribution of neutralization potency for mAbs isolated from both donors (**Fig. 3A and 3B**). 11.7% and 7.4% of the expressed F-binding mAbs (ELISA) from Donors 2.3 and 4.2, respectively, failed to appreciably neutralize the virus. Conversely, 50.7% and 55.5% of antibodies from Donors 2.3 and 4.2, respectively, demonstrated moderate (0.1–1 µg/mL) or strong (<0.1 µg/mL) neutralization. To further understand how neutralization potency related to binding specificity, we evaluated mAb neutralization in terms of preF and postF ELISA specificity (**Fig. 3C, 3D**). For both donors, >75% of hMPV F-binding mAbs were able to bind both pre- and postfusion F to some degree (**Fig. 3C**). Donor 2.3 had a greater frequency of preF preferring mAbs (*i.e.*, 10x or greater ELISA titer to preF than postF), and all mAbs that were entirely preF specific were also isolated from this Donor. Donor 4.2 had a higher frequency of mAbs with comparable binding to both preF and postF. There was no obvious segregation of neutralization with preF or postF binding preference (**Fig. 3D**). Interestingly, several of the isolated mAbs from both donors demonstrated postF specificity despite being sorted with a preF probe. This finding likely represents some contaminating postF either in the original protein preparation or that arises due to tagging with the fluorescent marker.

**Figure 3.**
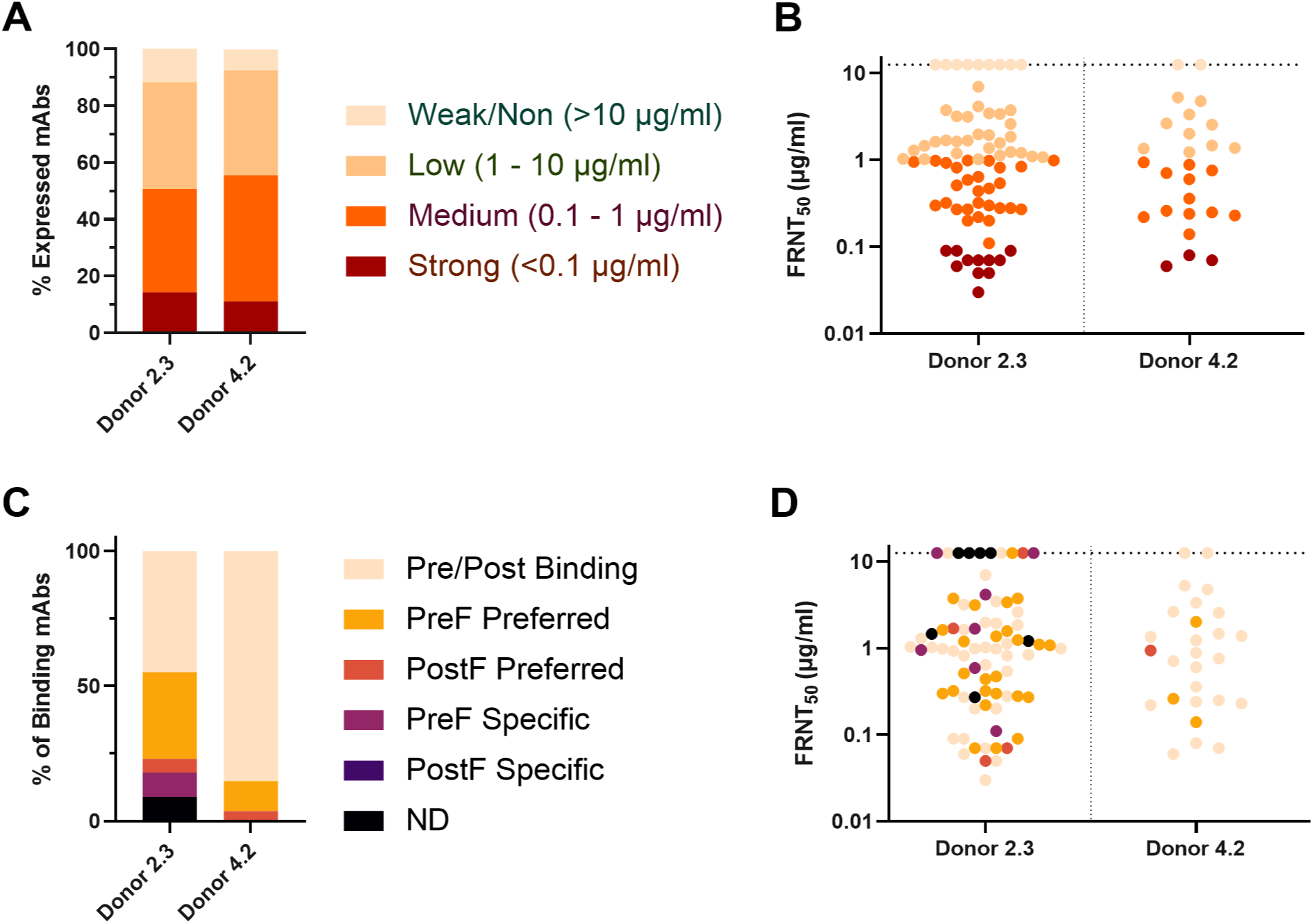
Antibody preF/postF specificity and neutralization potency by donor. All percentages were calculated based on the number of F-binding mAbs from the indicated donor. **A.** Percentage of expressed mAbs from each donor with the indicated potency. **B.** Neutralization potency of individual F-binding mAbs from both donors. **C.** Percentage of antibodies binding to hMPV preF or postF by ELISA. ND indicates antibodies that did not have reportable values for one of the assays required for analysis. The antibodies were defined as “Preferred” if they demonstrated 10-fold or greater ELISA binding titer to one of the protein conformations. The antibodies were defined as “Specific” if the observed binding titer was <333 ng/mL for one format and >333 ng/mL for the other format. **D.** Neutralization potency of mAbs, with colors reflecting preF/postF specificity.

### Epitope binning by competition binding

To map the binding sites of the isolated antibodies we conducted a competition assay using biolayer interferometry. Briefly, a prefusion-stabilized hMPV F construct was loaded onto biosensors and saturated with known antibodies. The saturated antigen was then used to test for binding of the isolated hMPV F-specific antibodies. Antibodies DS7 (site I), 338 (site II), MPE8 (Site III), and 101F (site IV) were used as the saturating antibodies to map the indicated antigenic sites (**Figs. 4, S1**). We found that 59 of the 106 F binding antibodies (ELISA) competed with one or more of the four aforementioned antibodies for binding to hMPV preF, whereas 37 of the 106 were non-competitive (**Figs. S1, S2**). These 37 non-competitive antibodies were then competed against each other, resulting in the identification of four additional epitope bins (A, B, C, and D). From the cryo-EM studies described below, Bins B and C were found to be equivalent to the prefusion-specific sites V and Ø, respectively. Bin A was found to be located within the trimer interface, similar to the previously described antibody MPV458, which binds near the apex of the F protein (**Fig. S3**) (Huang et al., 2020). Bin D antibodies were unable to bind to a preF construct that contained an engineered disulfide bond near the C-terminus (residues 365 and 463) – referred to in Methods as DS-CavEs2. Therefore, Bin D was determined to have a membrane-proximal epitope near the base of the preF conformation (**Figs. 4, S4**).

**Figure 4.**
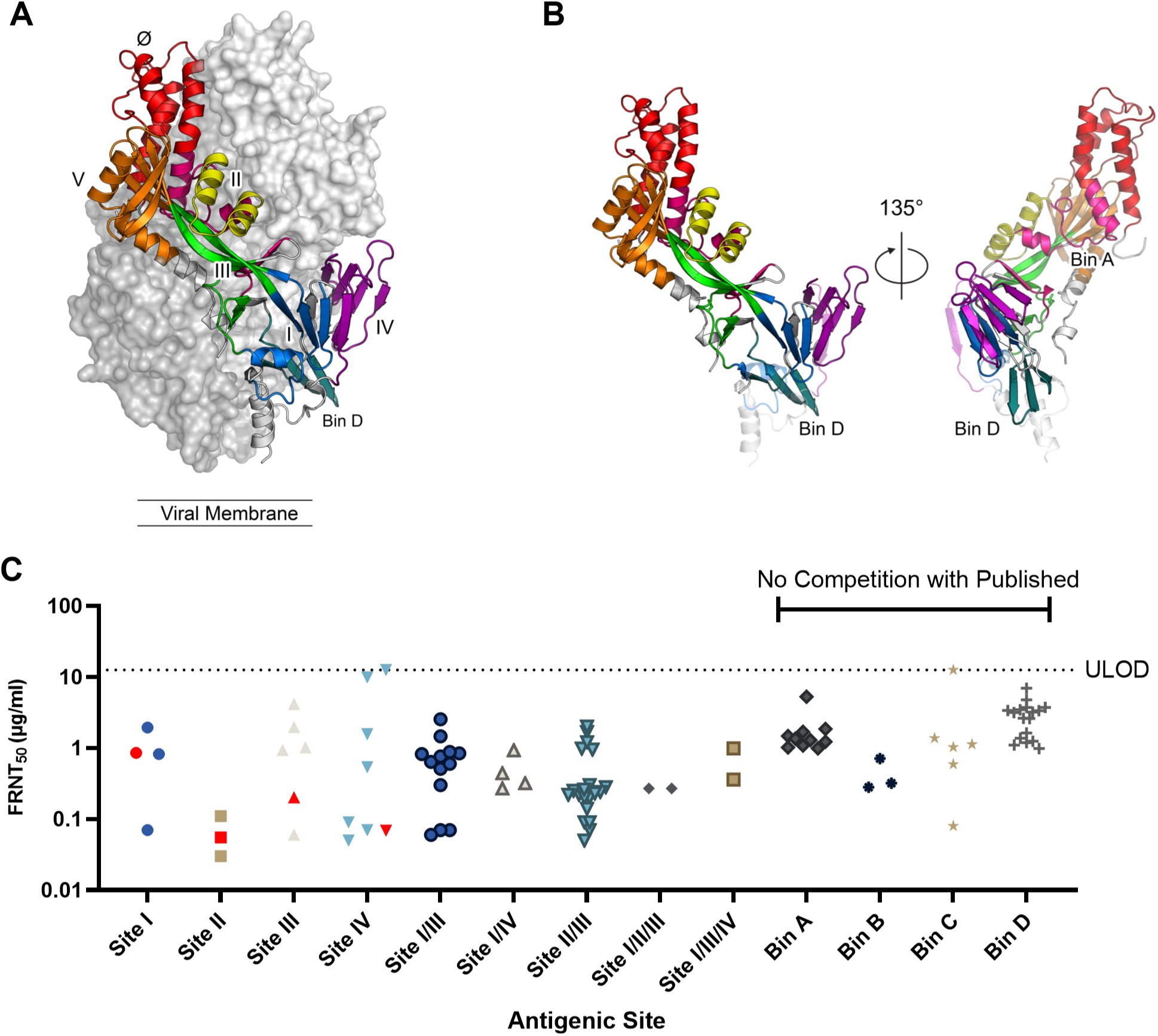
hMPV F antigenic sites and hMPV strain A1 virus neutralization, sorted by epitope bin. **A.** Structure of an hMPV preF trimer (PDB: 5WB0) with a single protomer shown as a ribbon colored to represent the antigenic sites. **B.** A single hMPV preF protomer is shown to better visualize Bin A and Bin D antigenic sites. Colored as in **A. C.** Virus neutralization titer for each mAb is shown according to the epitope bins defined in **A**, **B**. The neut titers are reported as the antibody concentration resulting in a 50% reduction in the number of focus-forming units in each well (FRNT_50_). Previously published antibodies that bind to antigenic sites I (DS7), II (338), III (MPE8) and IV (101F) are shown in red.

The majority of the isolated antibodies could be placed into just 3 of the 13 bins: Sites I/III overlapping, Sites II/III overlapping, and Bin D. Interestingly, we did not observe an obvious segregation of potency to specific antigenic sites. Most bins had at least a few neutralizing antibodies with IC_50_ <1 µg/mL, with the exception of Bin A and Bin D. This is in contrast to what has been observed for RSV F, where the most potent antibodies primarily target the prefusion-specific sites Ø and V, as well as antigenic site IV which is present on both the preF and postF conformations (Gilman et al., 2016). For hMPV F, only 1 of the 9 antibodies targeting sites Ø and V had an IC_50_ < 0.1 µg/mL. The majority (12/14) of the antibodies with IC_50_ < 0.1 µg/mL competed with antibodies 338, MPE8, and/or 101F, suggesting that the majority of the antibody response to hMPV F as a result of natural infection does not target prefusion-specific epitopes.

### Cryo-EM studies of two antibodies from unknown competition bins

We determined a cryo-EM structure of the SAN27-14 antigen-binding fragment (Fab) from Bin B bound to hMPV F to a global resolution of 3.1 Å. The MPE8 Fab, which has a quaternary epitope, was included in the complex to keep hMPV F in a compact trimer on the grids. Local refinement after subtracting the constant regions of both Fabs led to a better-resolved map of the Fab interfaces for model building (**Fig. 5A**). The structure revealed that SAN27-14 binds to antigenic site V—in proximity to MPE8 (site III) (Wen et al., 2017)—exclusively via its heavy chain, which buries 726 Å^2^ of surface area (**Fig. 5B**). The 15-amino-acid-long CDRH3 loop, stabilized by a disulfide bond, packs against the hMPV F surface formed by α3 and β3 from the F_1_ subunit. An *N*-linked glycan attached to Asn96 of the CDRH3 forms a hydrogen bond with the sidechain of Asn139 on F (**Fig. 5C**). In addition, the sidechain of CDRH1 Tyr33 forms a hydrogen bond with the sidechain of hMPV F Gln127. hMPV F Leu130, residing in a short loop connecting α2 and α3, is surrounded by Tyr33 and Tyr53 in the CDRH1 and CDRH2 loops, respectively (**Fig. 5D**). Notably, the CDRH2 and subsequent framework region reach toward antigenic site IV on the neighboring hMPV F protomer, establishing a quaternary contact via three salt bridges (**Fig. 5E, F**). These interactions involve hMPV F residues Lys429 and Glu431 at the hinge region preceding β22, which needs to flip 180° to adopt the postfusion conformation. Collectively, these data demonstrate that SAN27-14 binds to preF at regions in site V and site IV that refold during the pre-to-postfusion transition. These findings suggest that SAN27-14 neutralizes hMPV by preventing refolding of hMPV F, and thus inhibiting fusion of the viral and host-cell membranes.

**Figure 5.**
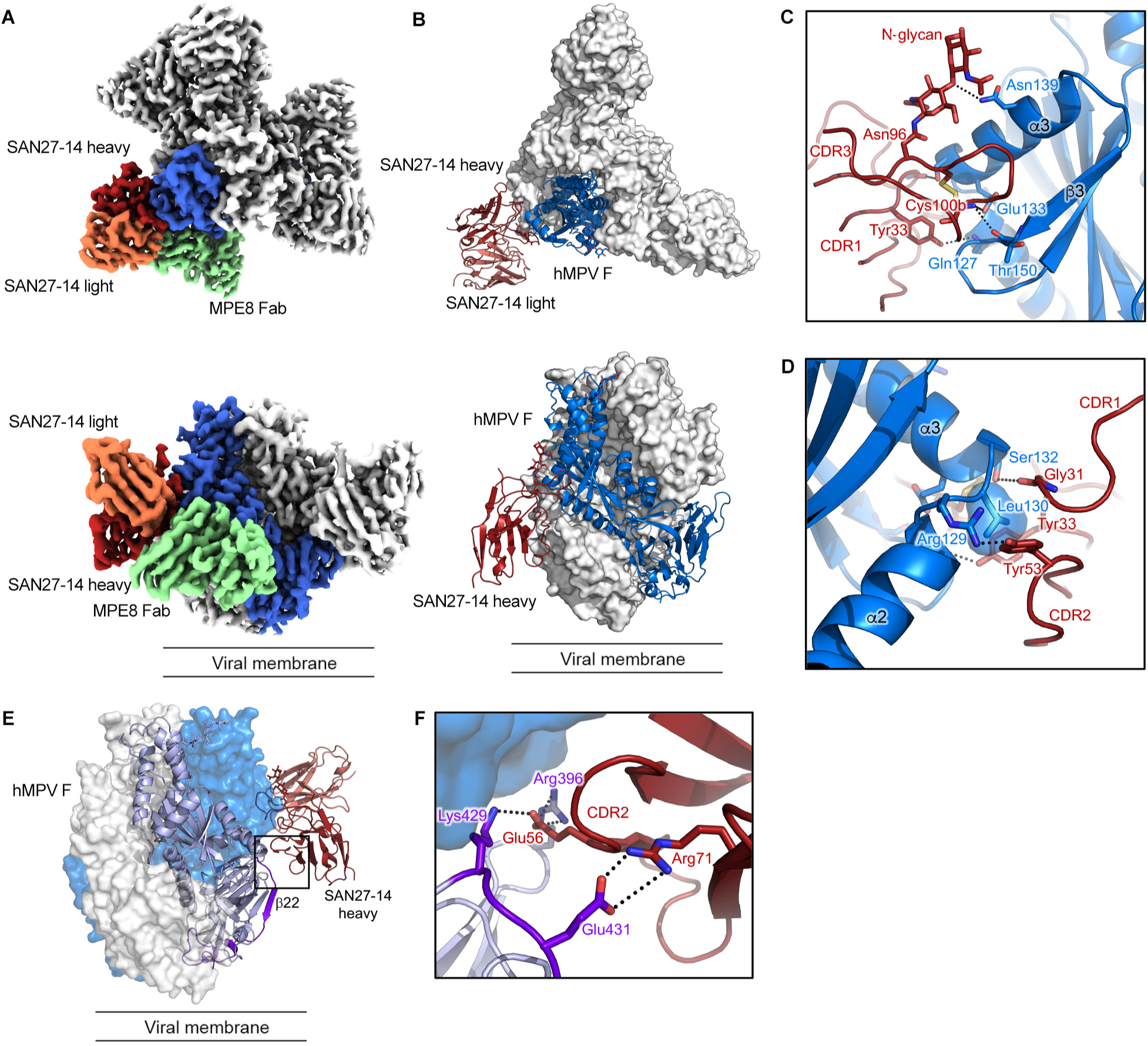
SAN27-14 Fab binds prefusion hMPV F at site V. **A.** Cryo-EM map of the SAN27-14 complex with MPE8-bound hMPV F shown in top and side views. A single protomer of the F trimer and its Fabs are colored (F: blue; SAN27-14 heavy and light chains: firebrick red and coral; MPE8 Fab: green). Constant domains of the Fabs have been removed for clarity. **B.** The model of top and side views of the protein complex with a single protomer and its Fabs shown as ribbons in the same color scheme as **A**. Remaining protomers and Fabs are depicted as a white surface. The MPE8 Fabs are removed for clarity. Only the heavy chain of the SAN27-14 is shown in the side view to emphasize the heavy chain-exclusive interaction. The F interacting N-glycan of Asn96 from SAN27-14 heavy chain is shown as sticks. **C, D.** Zoomed view of the SAN27-14 interface with hMPV F. Important residues are depicted as sticks. Polar interactions are depicted as dotted lines. **E, F.** Side and zoomed views of SAN27-14 bound to a quaternary epitope across two F protomers. The F protomer that forms the main interactions (**A-D**) with SAN27-14 is shown as a blue surface, and the other F protomer that forms three salt bridges (dotted lines) with SAN27-14 heavy chain is depicted as purple ribbons.

We also determined a cryo-EM structure of SAN32-2 Fab (Bin C) bound to hMPV F complexed with MPE8 Fab to a global resolution of 3.1 Å (**Fig. 6A, Table S5**). Local refinement focused on the variable regions of the SAN32-2 Fabs and the membrane-distal half of the F trimer led to a better-resolved map of the Fab interfaces for model building. The structure reveals that SAN32-2 binds to antigenic site Ø, with its heavy and light chains burying 359 Å^2^ and 394 Å^2^ of surface area, respectively, on hMPV F (**Fig. 6B**). CDRH2 residues Asp52 and Asp54 form salt bridges with hMPV F Lys179, which resides in the loop between α4 and α5 that rearranges into an extended helix during the preF-to-postF transition (**Fig. 6C**). CDRH2 residue Arg50 and CDRH3 residue Ser98 form hydrogen bonds to Asn180 in this loop as well. The SAN32-2 light chain binds to hMPV F through a network of hydrogen bonds formed between CDRL1 Tyr32 and hMPV F Asp72 as well as between multiple CDRL2 residues and Arg79 of hMPV F (**Fig. 6D**). Additionally, CDRL3 Asp92 forms a salt bridge with hMPV F Lys68 in the α1 helix.

**Figure 6.**
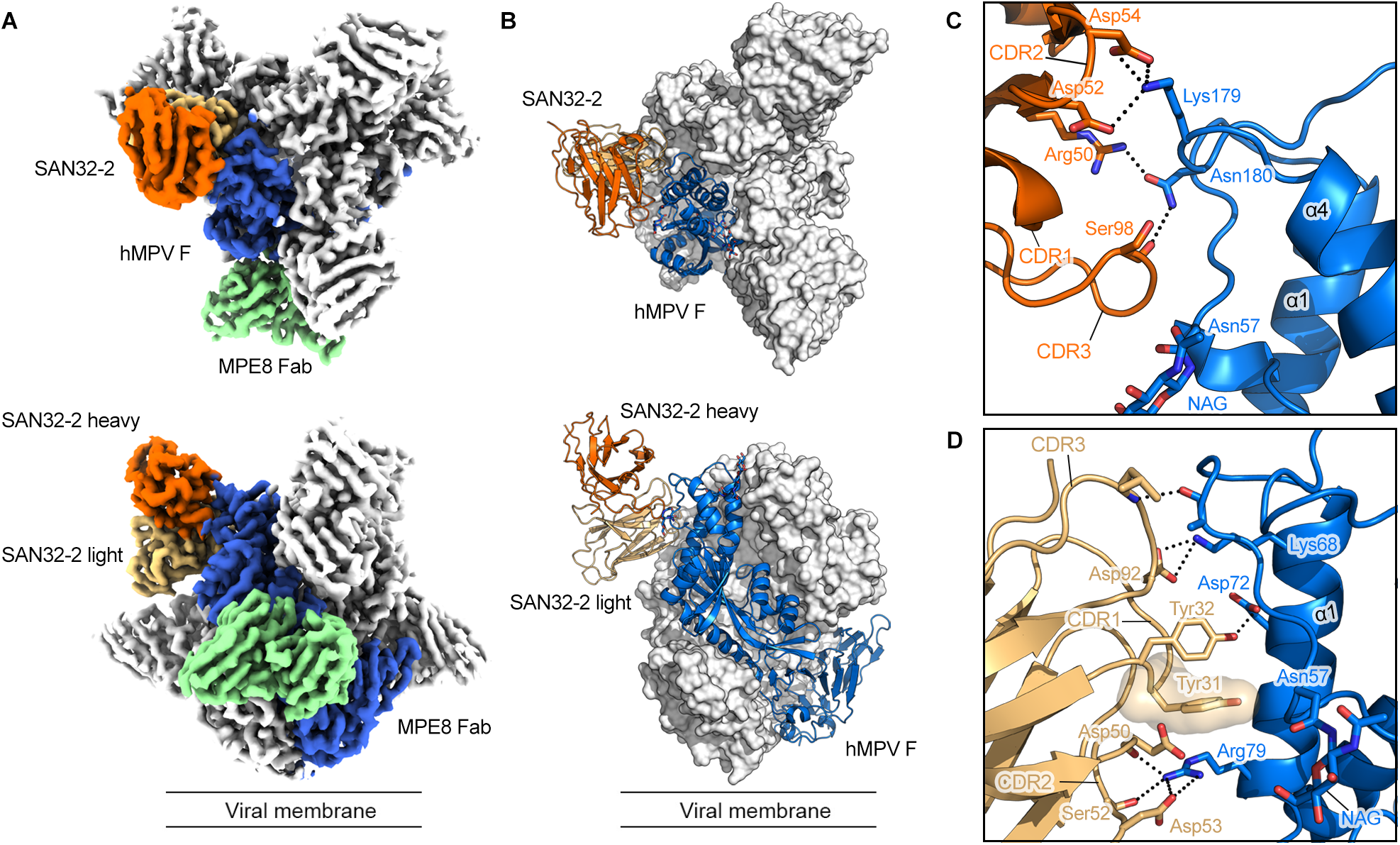
SAN32-2 Fab binds prefusion hMPV F at site Ø. **A,** (top) View from above the SAN32-2 protein complex cryo-EM map. A single protomer of the trimer and its Fabs are colored (F: blue; SAN32-2: orange; MPE8: green). Constant domains of the Fabs have been removed for clarity. (bottom) Side view of the cryo-EM map. **B.** (top) View from above the protein complex with the model of a single protomer and its Fabs shown as colored ribbons(F: blue; SAN32-2 heavy chain: dark orange; SAN32-2 light chain: light orange). Remaining protomers and Fabs are depicted as a surface representation colored white. The MPE8 Fab and constant domains have been removed for clarity. (bottom) Side view of the protein complex colored the same as the top view. Only the variable domain of the SAN32-2 model is shown, all other fabs removed for clarity. **C, D.** Zoomed in view of the SAN32-2 interface with hMPV F. Important residues are shown as sticks. Hydrogen bond and salt bridge are depicted as dotted lines.

Compared to RSV F, hMPV F has an additional glycosylation site within antigenic site Ø at Asn172 in α4. We had previously suggested that this would create a glycan shield at the apex of hMPV preF and reduce elicitation of site Ø-directed antibodies (Battles et al., 2017). Indeed, only six antibodies were identified as belonging to this competition bin. SAN32-2 is able to evade the glycan shield by binding toward the α1 helix of F_2_ with a horizontal angle of approach (**Fig. 6B**). SAN32-2 fits between the Asn57 glycan of the protomer to which it is bound and the Asn172 glycan from the neighboring protomer. The SAN32-2 structure thus defines a site Ø epitope on hMPV F and reveals how a neutralizing antibody can evade the glycan shield.

## DISCUSSION

Only the fusion protein elicits neutralizing antibodies as a result of hMPV infection (Ryder et al., 2010; Skiadopoulos et al., 2006). It is therefore important to understand the antibody response to hMPV F and how its epitopes correlate with neutralization. Although previous studies generated hMPV F-specific antibodies (Bar-Peled et al., 2019; Huang et al., 2020; Mas et al., 2016; Ulbrandt et al., 2008; Ulbrandt et al., 2006; Wen et al., 2012; Williams et al., 2007; Wu et al., 2007), the mapping of the epitope landscape remained largely incomplete. Initial reports suggested that the majority of the neutralizing response was directed toward epitopes shared on the preF and postF conformations (Battles et al., 2017; Pilaev et al., 2020). In the current study, we successfully utilized a prefusion-stabilized hMPV F probe to isolate and sequence the immunoglobulin heavy and light chain variable regions from circulating B cells of two elderly donors who were naturally infected with hMPV, likely multiple times throughout their lives. We then characterized the preF-binding antibodies using biolayer interferometry and cryo-EM to comprehensively map the antigenic sites of hMPV preF. Our results indicate a broad surface coverage with representative mAbs binding to all six RSV F-analogous antigenic sites (Ø, I, II, III, IV, and V), as well as two novel antigenic sites which have not been previously described (Bin A and Bin D).

Unlike what has been observed for the closely related RSV (reviewed in (Ruckwardt et al., 2019)), targeting of preF-specific epitopes V and Ø does not appear to be critical to achieve potent neutralization of hMPV. The antigenic sites for the potently neutralizing antibodies described here are diverse and there is no clear segregation to dominant sites. Our panel of antibodies included 14 mAbs with a FRNT_50_ titer ≤0.1 µg/mL, and these 14 mAbs were found to bind to Sites Ø, I, II, III, and IV, as well as hybrid sites II/III and I/III. For RSV, site V has been shown to be the target of very potent neutralizing antibodies (Gilman et al., 2016); however, the three site V antibodies isolated in this study had moderate to low FRNT_50_ titers between 0.1 and 1 µg/mL.

Surprisingly, we did not isolate a single antibody that was cross-reactive with RSV F. Prior work has led to the discovery of RSV F/hMPV F cross-reactive mAbs such as MPE8 (Site III, (Corti et al., 2013)), 101F (Site IV,(Wu et al., 2007)), and M1C7 (Site V,(Xiao et al., 2019)). In our work, we isolated five site III-specific mAbs, eight site IV-specific mAbs, and three site V-specific mAbs that were all hMPV specific. All of these had distinct variable gene usage and no obvious sequence convergence with previously published cross-reactive antibodies, which likely explains the absence of RSV cross-reactivity (Xiao et al., 2019). In addition, a number of previously reported cross-reactive antibodies utilized the VH3-21 heavy chain germline and the VL1-40 light chain germline (Corti et al., 2013; Gilman et al., 2016; Xiao et al., 2019). We did observe this pairing in several of the sequences from one subject, but they have not yet been expressed and characterized.

We observed diverse heavy chain germline gene usage amongst the preF-sorted B cells, with several interesting deviations from a published, unsorted B cell repertoire (Boyd et al., 2010). In particular, we noticed a striking enrichment of IGHV1-2 and IGHV3-30 in the preF-sorted B cells from these donors. The 39 antibodies we expressed to represent these germline variable regions were varied in the antigenic sites recognized as well as neutralization potency, though there was a relatively high frequency of site II/III-specific antibodies from the IGHV3-30 heavy chain germline (7 of 23), and Bin D-specific antibodies from the IGHV1-2 heavy chain germline (7 of 16). Though these data are only derived from 2 donors, it seems likely that naïve B cells carrying these heavy chain variable regions are disproportionately enriched by hMPV infection. Such information can potentially be leveraged to guide antigen design as well as future antibody discovery campaigns.

Notably, we were able to identify two novel hMPV F antigenic sites (Bin A and Bin D). Bin A antibodies did not compete with any of the antibodies used for competition, nor did they compete with our site V and site Ø antibodies. However, a subset within Bin A was shown to partially compete with the previously described trimer-interface-binding antibody MPV458, which binds a linear epitope comprising amino acids 66–87 on the α1 helix (**Fig. S3**) (Huang et al., 2020). The existence of an antigenic site that resides at least partially within the trimer interface suggests, as previously speculated (Huang et al., 2020), that uncleaved hMPV F on the surface of the virus is monomeric or that the trimer transiently ‘breathes’ to expose this site, as has been demonstrated for influenza HA (Bajic et al., 2019; Bangaru et al., 2019; Watanabe et al., 2019) and RSV F (Gilman et al., 2019). Furthermore, the largest competition bin in our study (Bin D) is near the membrane-proximal base of the protein (**Fig. S4**). Nanobodies recognizing a similar region on RSV F have recently been described (Xun et al., 2021) and are potent neutralizers, whereas Bin D neutralizing antibodies in our study are rare, with only 1 of the 17 having strong neutralization potency (**Fig. S1**). Further investigation into Bin A and D antibodies may ultimately uncover more potently neutralizing antibodies.

The results described here should facilitate development of an effective hMPV vaccine by providing a better understanding of the humoral response to hMPV F-elicited natural infection. Additionally, the antibodies from this study will be useful reagents for future investigation of hMPV F, and the most potently neutralizing antibodies may prove to be useful as therapeutic candidates.

## ACKNOWLEDGMENTS

We thank: members of the McLellan Laboratory for providing helpful comments on the manuscript; Dr. Sasha Dickinson at the Sauer Structural Biology Laboratory at UT Austin for assistance with cryo-EM data collection; Dr. John Ludes-Meyers and Dr. Kaci Erwin for mammalian protein expression assistance; Dr. Douglas Zhang at AbHelix, LLC for his support in NGS data acquisition and analytics. We acknowledge the University of Texas College of Natural Sciences and award RR160023 of the Cancer Prevention and Research Institute of Texas for support of the EM facility at the University of Texas at Austin. This work was funded in part by Welch Foundation grant number F-0003-19620604 (J.S.M.).

## AUTHOR CONTRIBUTIONS

Conceptualization, J.M.D., J.S.M., T.M.F., L.Z.; Investigation and visualization, S.A.R., G.B., C.L.H., E.C., J.N.R.T., C.S., C.A.B., A.K., J.K.; Writing – Original Draft, S.A.R., G.B., C.L.H., E.C., C.A.B.; Writing – Reviewing & Editing, S.A.R., G.B., C.L.H., E.C., J.N.R.T., C.S., C.A.B., S.M., L.Z., J.M.D., J.S.M.; Supervision, R.G., S.M., D.C., T.M.F., J.M.D., J.S.M.

## DECLARATION OF INTERESTS

Gurpreet Brar, Emilie Chautard, Jennifer N. Rainho-Tomko, Christine A. Bricault, Rachel Groppo, Sophia Mundle, Linong Zhang, Danilo Casimiro, Tong-Ming Fu, and Joshua M. DiNapoli are all current or former employees of Sanofi Pasteur and may hold stock in Sanofi company.

## METHODS

### Patient Selection

These studies were conducted with volunteer serum samples obtained through Sanofi Pasteur VaxDesign (Orlando, FL). The collections and study protocol were reviewed and approved by Chesapeake Research Review Inc (Columbia, Maryland) under IRB 0906009. Sera utilized in these experiments were collected from healthy elderly individuals, ages 59–76. Patient samples were aliquoted and stored in liquid nitrogen at -130 °C until use. Donor plasma and peripheral blood mononuclear cells (PBMCs) were cryopreserved and stored for an extended period of time in vapor phase nitrogen tanks.

### Recombinant protein production

An hMPV F A1 gene encoding residues 1–490 from strain NL/1/00 with the putative F cleavage site changed to a furin cleavage site (ENPRQSR -> ENPRRRR), as well as substitution of A185P and G294E, was cloned into mammalian expression vector pαH upstream of a C-terminal “GGGS” linker sequence followed by the T4 fibritin trimerization motif “foldon” (Efimov et al., 1994; Miroshnikov et al., 1998), an HRV3C protease cleavage site, an 8xHis tag, and a Strep-TagII (Battles et al., 2017). Gibson assembly was used to introduce the prefusion-stabilizing disulfide bond A113C/A339C (Stewart-Jones et al., 2021) and an H368N (Schowalter et al., 2009) substitution. This protein is referred to as 115-BV-DS. An hMPV F B2 (strain TN99-419) version of this construct was synthesized by GenScript. Protein expression was carried out using FreeStyle 293F cells (ThermoFisher) transiently co-transfected at a 4:1 ratio of hMPV F and furin-expressing plasmids using polyethylenimine (PEI). A final concentration of 5 µM kifunensine and 0.1% (v/v) Pluronic F-68 (Gibco) were added to the cells three hours after transfection. After six days, protein was purified via Strep-Tactin Sepharose resin (IBA) from cell supernatants that had been filtered and buffer-exchanged into PBS by tangential flow filtration. Protein was further purified by size-exclusion chromatography using a Superose 6 Increase 10/300 column (GE Healthcare) in 2 mM Tris pH 8.0, 200 mM NaCl, and 0.02% NaN_3_ running buffer.

The prefusion hMPV F construct DS-CavEs2 (hMPV F A1 NL/1/00, residues 1-490), which was used for structural studies, includes the G294E, A185P, and furin cleavage site substitutions as described above as well as disulfides at L110C/N322C, T127C/N153C, A140C/A147C, T365C/V463C and additional substitutions of L219K, V231I, and E453Q (*manuscript under review*). Expression and purification were as described above.

Postfusion hMPV F A1 and B2 constructs were produced as previously described (Mas et al., 2016). Transient transfection of FreeStyle 293F (Thermo Fisher) cells was as described above. After six days, cell supernatant was buffer exchanged into 20 mM Tris pH 8.0, 20 mM imidazole pH 8.0, 300 mM NaCl and passed over Ni-NTA resin. After a wash step with 20 mM Tris pH 8.0, 50 mM imidazole, 300 mM NaCl, protein was eluted from resin with 20 mM Tris, pH 8.0, 250 mM imidazole pH 8.0, 300 mM NaCl. Additional purification was then carried out by size-exclusion chromatography using a Superose 6 Increase 10/300 column (GE Healthcare) with 2 mM Tris pH 8.0, 200 mM NaCl, and 0.02% NaN_3_ running buffer. After purification, protein was heated at 70 °C for 10 minutes to convert any residual prefusion F to postfusion F.

### Sorting probe generation

To generate antigen-specific probes for sorting, trimerized hMPV A1 prefusion-stabilized F protein (115-BV-DS, described above) was singly biotinylated with the Thermo EZ-Link Micro NHS PEG4 Biotinylation Kit (Cat: 21955) before being conjugated to streptavidin-PE or -APC (Gilman et al., 2016; Goodwin et al., 2018). Streptavidin-PE and -APC were conjugated to the biotinylated hMPV A prefusion F proteins at a 4:1 molar ratio, respectively. Tetramers were made fresh for each experimental run.

### Sorting

B cells were enriched from human PBMCs using EasySep Human Pan-B Cell Enrichment Kit (StemCell Technologies) before being incubated with Fc block (BD Biosciences) for 15 mins at 4 °C. Cells were then stained with Aqua Live/Dead FITC (Invitrogen), CD3 PerCP/Cy5.5 (BD Bioscience), CD8a PerCP/Cy5.5 (BD Bioscience), CD14 PerCP/Cy5.5 (BD Bioscience), SA PerCP/Cy5.5 (BD Bioscience), CD19 PE/Cy7 (BD Bioscience), CD20 PE/Cy7 (BD Bioscience), IgM BV421 (Biolegend), SA/PE/preF hMPV (PE; 200 nM), SA/APC/preF hMPV (APC; 200 nM) for 30 min at 4°C. Live, singlet cells that were CD3-, CD8a-, CD14-, SA-, CD19+, CD20+, IgM-, SA/PE/preF hMPV+, SA/APC/preF hMPV+ B cells were sorted using a Sony SH800S Cell Sorter.

### Single-cell library construction

Antigen-sorted cells were loaded into Chromium A chips (10x Genomics) and converted into gel beads-in-emulsion (GEMs) according to the manufacturer’s guidelines using the Single Cell 5′ Bead Kit (10X Genomics) and the Chromium Single Cell A Chip Kit (10X Genomics). Once the GEMs were generated, samples underwent a reverse transcription reaction with reagents provided by the Chromium Single Cell 5′ Library Kit (10X Genomics). GEM-RT products were subsequently amplified using human BCR primers and cleaned up through SPRIselect beads (Beckman Coulter). Next, products were further prepared using the Single Cell V(D)J Enrichment Kit, Human B Cell Kit (10X Genomics) and the quality and quantitation of the products were monitored using the High Sensitivity DNA Kit (Agilent Technologies). Libraries were sequenced on an Illumina sequencer with Illumina adaptor primers found in the Chromium i7 Multiplex Kit.

### Single cell V(D)J data analysis

A quality check of raw data sequences was initially performed with FastQC. A first analysis was then performed to extract V(D)J sequence contigs from FASTQ files on samples 2.3 and 4.2 using the Cell Ranger VDJ pipeline from 10x Genomics, Cell Ranger versions 3.0.2 and 3.1.0 respectively (Zheng et al., 2017). This initial analysis was used to select the antibodies to be expressed for each sample. Sequence contigs not annotated as productive, or with no raw clonotype identifier or C gene call (constant region) were dropped. Sequences with more than one defined sequence per cell identifier for heavy and light chains were considered doublets and dropped. Paired heavy chain and light chain sequences were then analyzed using the Geneious software to select the candidates for expression. Sequences were selected for expression based on coverage of diverse lineages and clonotypes, as well as based on sequence features deemed of interest.

A second in-depth analysis was subsequently performed using an updated and upgraded analysis pipeline. The raw data were first processed using the Cell Ranger VDJ 6.0 pipeline from 10x Genomics. Assembled contigs from Cell Ranger were then analyzed using a custom bioinformatics pipeline based on the open-source Immcantation package (https://immcantation.readthedocs.io/en/stable/). Contigs were re-aligned with IgBLAST using the Immcantation changeo-10x superscript based on Change-O (v1.0.2) (Gupta et al., 2015). The superscript was used to run the AssignGenes.py IgBLAST wrapper with Ig chains on IMGT database (2021.03.26), MakeDb.py to process IgBLAST output and ParseDb.py to select IGH and IGK/IGL loci. Productive vs non-productive sequences were split, but no clonal assignment was performed at this step. Heavy and light chain sequences were merged using a custom R script and R.3.6.3. Contig sequences with no amino-acid sequences for junctions were dropped. We also dropped sequences with more than one defined sequence per cell identifier for heavy and light chains, which were considered doublets. Light chains were merged with Heavy chains based on their cell identifier, and all light chains with no corresponding heavy chain were dropped. Custom R scripts were used to annotate the sequences. Only productive sequences, with no stop codons, and with VJ in frame, were kept. Amino acid properties were calculated with the aminoAcidProperties function from alakazam R package (v1.0.2). The observed number of mutations was defined with the observedMutations function from shazam R package (v1.0.2) (Gupta et al., 2015). The superscript changeo-clone.sh was used to define clones (Change-O v1.0.2 script DefineClones.py) and to assign germlines (CreateGermlines.py). BCR sequences that have the same heavy chain V and J genes and same junction length were grouped. Sequences with similar CDR3 junction regions were then clustered with a single-linkage hierarchical clustering into clonal groups, using a defined length-normalized nucleotide Hamming distance of 0.15. After the germline assignment step, a custom R script was used to define clones, clonotypes and lineages. Clones were defined as sequences that are identical at the nucleotide level for V and J genes as well as the CDR3 nucleotide sequence. Clonotypes were defined as sequences that have the same V and J gene call and the same CDR3 amino acid sequence. Lineages were defined as having the same V and J gene call, as well as a CDR3 within a normalized nucleotide Hamming sequence of 0.15. This lineage definition is equivalent to the clonal group definition for the Immcantation package.

The expressed antibodies, which were selected from the initial Cell Ranger v3 analysis, were reconciled with Cell Ranger 6.0 results using the cell barcode information. As some assembled sequences were considered high-confidence cellular contigs only in Cell Ranger 6.0 (e.g., rearrangements not productive or not cell-associated), some of the antibodies selected for expression (19 contigs out of 100 for sample 2.3 and 2 out of 36 for sample 4.2) were not processed by the Immcantation pipeline to identify lineages and germlines sequences. Apart from these sequences, all contigs from Cell Ranger 3 and 6 had the same CDR3 amino acid sequences. Several additional differences in the assembled contigs from Cell Ranger 3 vs Cell Ranger 6 were identified in the other regions (in 29 out of 79 contigs for sample 2.3, and 1 out of 34 for sample 4.2). The R package ggplot2 (Wickham, 2016) was used to create the plots for figure 2.

### Antibody and Fab Protein Production

Expression plasmids, full-length antibodies, and Fabs were synthesized, expressed, and purified by Sino Biological using their standard methods. Briefly, the variable regions of the heavy and light chain sequences obtained from the Cell Ranger output underwent sequence evaluation and non-canonical amino acids within the framework regions were replaced with canonical amino acids. Conserved leader signal sequences were used for both heavy and light chains. Heavy chain variable sequences were placed in a human IgG1 constant region backbone expression vector. Light chain variable sequences were placed into an expression vector containing a kappa or lambda light chain constant region, as determined by the original pairing of the mAb in the host. Genes were amplified by PCR and placed into expression vectors, which underwent sequence validation. Plasmids were transiently transfected into HEK293 cells in serum-free medium. After 6 days supernatant was harvested, spun down, and loaded onto Protein A affinity columns. Antibodies were assessed by SDS-PAGE. For preparation of Fabs, full-length antibodies were enzymatically cleaved with papain and Fab was purified by SEC-HPLC using an Agilent 1100 with a TSK G3000SWXL column.

### ELISA

Copper coated high binding capacity plates (Pierce) were coated with the hMPV prefusion F protein, referred to as 115-BV, at 1 μg/mL in PBS overnight at 4 °C. Plates were washed 3 times with PBS-Tween 0.1% before blocking with 1% BSA in PBS-Tween 0.1% for 1 h at ambient temperature. The plates then received a 3-fold, 8-point serial dilution of each antibody in blocking buffer, covering a dilution range of 1000 ng/mL to 0.4 ng/mL. Plates were washed 3 times after 1 h incubation at room temperature before adding 50 μL of 1:5000 Rabbit anti-human IgG (Jackson Immuno Research) to each well. Plates were incubated at room temperature for 1 h and washed 3 times. Plates were developed using Pierce 1-Step Ultra TMB-ELISA Substrate Solution for 0.1 h and stopped by TMB stop solution. Plates were read at 450 nm in a SpectraMax plate reader. Antibody titers were reported as the lowest antibody concentration resulting in an optical density >0.2.

### Viruses and Cells

HMPV strain CAN97-83 (A2) virus carrying a GFP reporter was obtained from ViraTree (Product Number: M121). CCL-81.2 Vero cells were cultured in OptiPRO (ThermoFisher) + 1% AA Antibiotic-Antimycotic (Gibco). PBMCs and serum were obtained from healthy donors.

### Microneutralization Assay

The neutralization activity of the antibodies was measured using CCL-81.2 Vero cells plated in Fluorobrite DMEM (ThermoFisher) + 1x Glutamax (ThermoFisher). Antibodies were 4-fold serially diluted (12.5 µg/ml – 0.012 µg/ml) and 75 µL of the diluted antibody was incubated with 75 µL WT hMPV-GFP1 virus at 32 °C for 1 hour. 50 µL antibody-virus mixture was added to the CCL-81.2 Vero cells plated in Fluorobrite DMEM + 1x Glutamax and incubated at 32 °C for 1 hour with rocking every 15 minutes. At the end of the incubation, 100 µL of Fluorobrite DMEM + 1x Glutamax was added to the wells and plates were incubated at 32 °C. Fluorescent cell counting of WT hMPV-GFP1 virus-infected Vero cells was performed using a Celigo microplate reader (Nexcelom Bioscience) after 48 hours of incubation. Neutralizing titer was reported as the lowest antibody concentration resulting in a 50% reduction of input virus focus forming units.

### Epitope Binning Assays

Competition binning was performed using an Octet RED96e system at 25 °C, with a shaking speed of 1,000 rpm using streptavidin (SA) biosensors (FortéBio). Biosensors were first equilibrated in running buffer (0.01 M HEPES, 150 mM NaCl, 0.05% Tween 20 and 1 mg/mL BSA) for 10 minutes before an initial 30 s baseline “sensor check” by dipping the biosensors into wells containing running buffer. Next, biosensors were loaded with antigen by immersion in wells containing 75 nM of hMPV preF A1 for 115 s followed by immersion in running buffer for 60 s to get a baseline. Then antigen loaded tips were immersed into wells containing primary mAb of either DS7 (400nM), 338 (200nM), MPE8 (200nM), 101F (200nM), or running buffer for 600 s, followed by another 60 s baseline step of buffer-only wells. Finally, saturated antigen was immersed into 100 nM of secondary mAb for 300 s and the final response signals were observed to determine competition with the primary antibody. After completion of the assay, biosensors were used two more times after being regenerated by three repeat sequences of dipping into 10 mM glycine pH 3.0 (60 s) then running buffer (60 s) before starting the next assay with the 30 s “sensor check” step. The antigen loading step for assays two and three were extended from 115 to 150 and 280 s, respectively. FortéBio software Octet® Data Analysis High Throughput v11.1 was used to generate an epitope binning competition matrix and the values were normalized to the binding signal of secondary antibody binding to the antigen alone. Data were then transferred to GraphPad Prism version 9.2.0 for data visualization. Antibodies were characterized as full competition (<0.33), partial competition (0.33–0.66), or no competition (>0.66) based on their relative binding value.

### Cryo-EM Sample Preparation and Data Collection

Purified hMPV preF DS-CavEs2 was combined with a 2-fold molar excess of MPE8 Fab and a 1.5-fold molar excess of either SAN32-2 or SAN27-14 Fab incubated in 2 mM Tris pH 8.0, 200 mM NaCl, and 0.02% NaN_3_ at room temperature for 10 minutes before being moved to ice. Just before freezing, 0.5 µL of 0.25% amphipol A8-35 was combined with 5 µL of sample and 4 µL was applied to a CF-1.2/1.3 grid that had been plasma cleaned for 60 seconds in a Solarus 950 plasma cleaner (Gatan) with a 4:1 ratio of O_2_/H_2_. Grids were plunge frozen using a Vitrobot Mark IV (Thermo Fisher). In a 10 °C, 100% humidity chamber, settings for sample blotting were 5 seconds of wait time followed by 4 seconds of blotting at a -2 force before plunging into liquid ethane. Using an FEI Titan Krios equipped with a K3 direct electron detector (Gatan), a single grid from each complex was imaged to collect a total of 3,492 micrographs for the SAN27-14 complex and 3,636 micrographs for the SAN32-2 complex. Data were collected at a 30° tilt with magnification of 29,000x corresponding to a calibrated pixel size of 0.81 Å/pix and a total dose of 70 e^-^/ Å^2^. Data collection statistics are in **Tables S4 and S5**.

### Cryo-EM data processing

Micrographs were corrected for gain reference and imported into cryoSPARC Live v3.2.0 for initial data processing: motion correction, defocus estimation, micrograph curation, particle picking and extraction, and particle curation through iterative streaming 2D class averaging. 2D averages were used to generate templates and template-based particle picking was carried out. Curated particles were exported to cryoSPARC v3.2 for further processing via rounds of 2D classification, *ab initio* reconstruction, heterogeneous refinement, and non-uniform homogenous refinement using C3 symmetry. Masking and particle subtraction were used for further non-uniform local refinement. Finally, both global and focused maps were sharpened using DeepEMhancer (Cianfrocco et al.; Sanchez-Garcia et al., 2021) and a combined focused map was generated in PHENIX. EM processing workflows are shown in **Figures S5 and S7**, and EM validation results are shown in **Figures S6 and S8**. For model building, an initial hMPV F model was generated from PDB ID: 5WB0 and a Fab model generated by SAbPred server (Dunbar et al., 2016) were used to initially dock into the cryoEM maps using UCSF ChimeraX (Pettersen et al., 2021). Models were built further and iteratively refined using a combination of C*oot* (Emsley et al., 2010), PHENIX (Liebschner et al., 2019), and ISOLDE (Croll, 2018). Model statistics are shown in **Tables S4 and S5**.

---

**Supplementary Figure 1.**
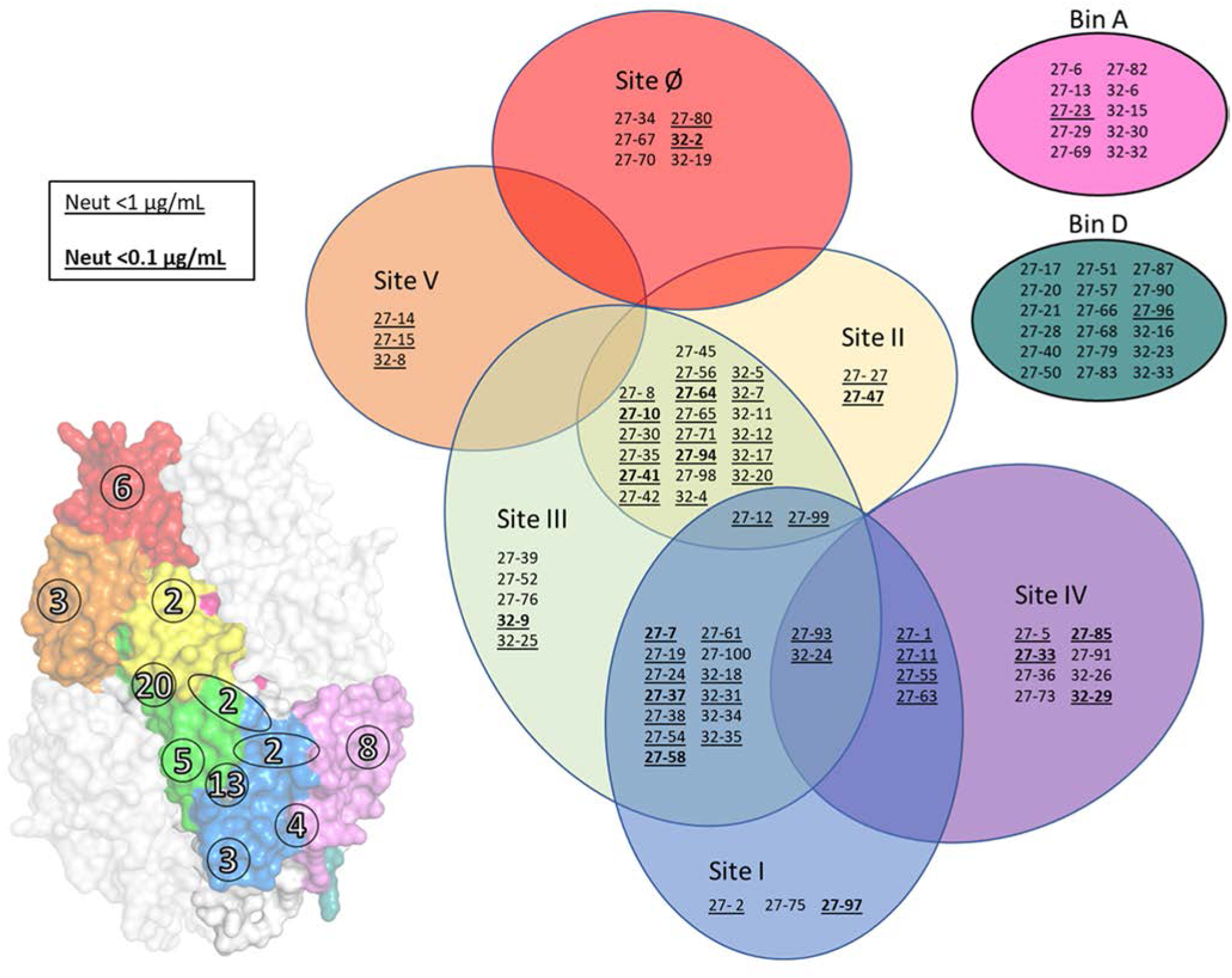
Antigenic landscape of isolated antibodies. An hMPV F trimer (left; PDB: 5WB0) is shown as a white surface rendering with a single protomer color coded to the antigenic sites . The number overlays represent the number of isolated antibodies binned to each respective site. A Venn diagram (right) of the antigenic sites, loosely organized as they would be on the protein surface, contains a list of the isolated antibodies binned to each specific antigenic site. Underlined antibodies are potent neutralizers, defined here as neutralizing titers < 1 µg/mL. Bold and underlined antibodies are very potent neutralizers, defined here as neutralizing titers < 0.1 µg/mL.

**Supplementary Figure 2.**
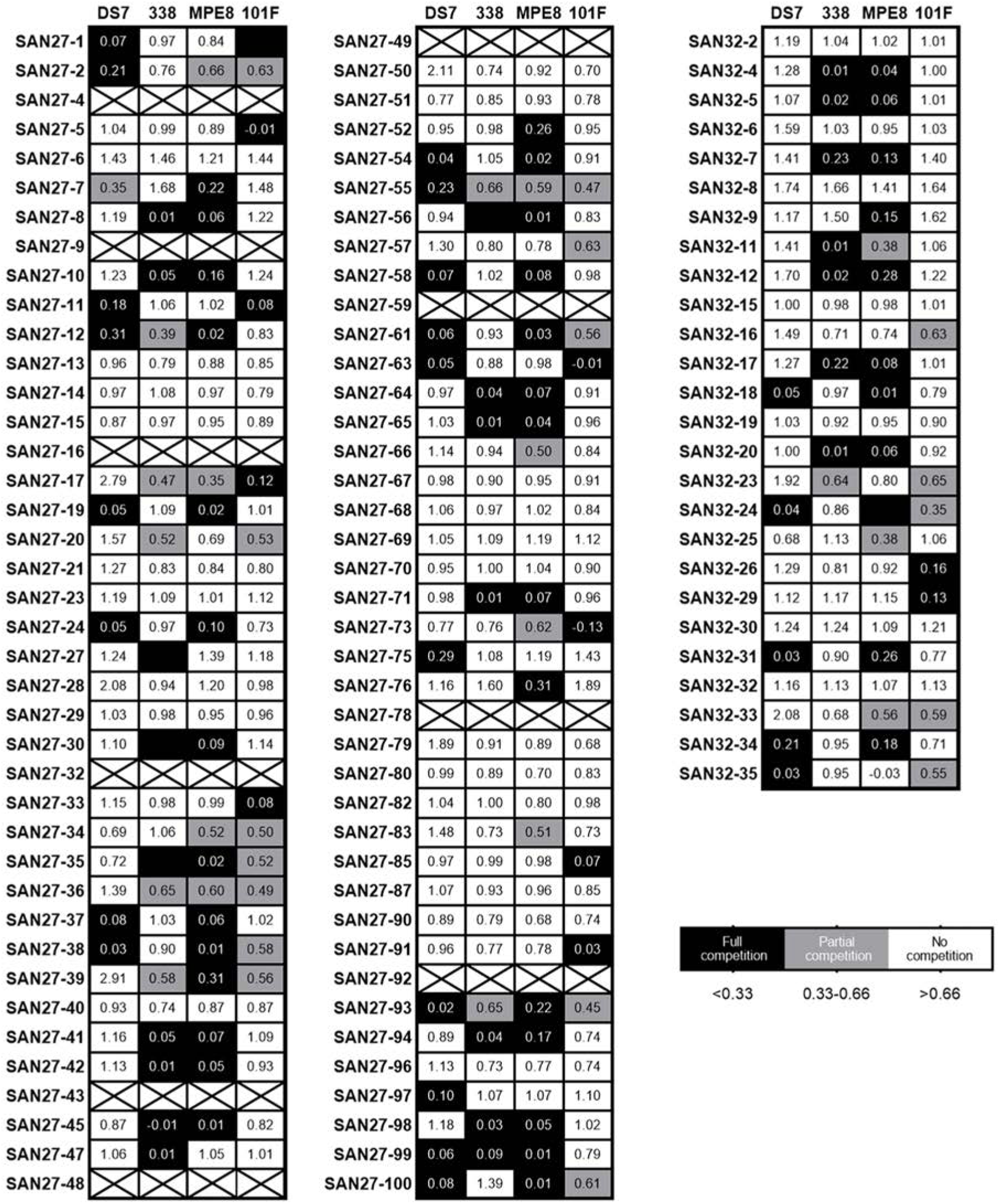
Competition matrix of isolated antibodies with published antibodies. The published antibodies [DS7(site I), 338(site II), MPE8(site III), and 101F(site IV)] used to saturate hMPV F are labeled at the top of the columns. Rows are labeled as the isolated antibodies. Numerical values within cells indicate normalized binding compared to the isolated antibodies’ binding to hMPV F. Rows with “X’ indicate antibodies that did not bind to hMPV F during the competition assay.

**Supplementary Figure 3.**
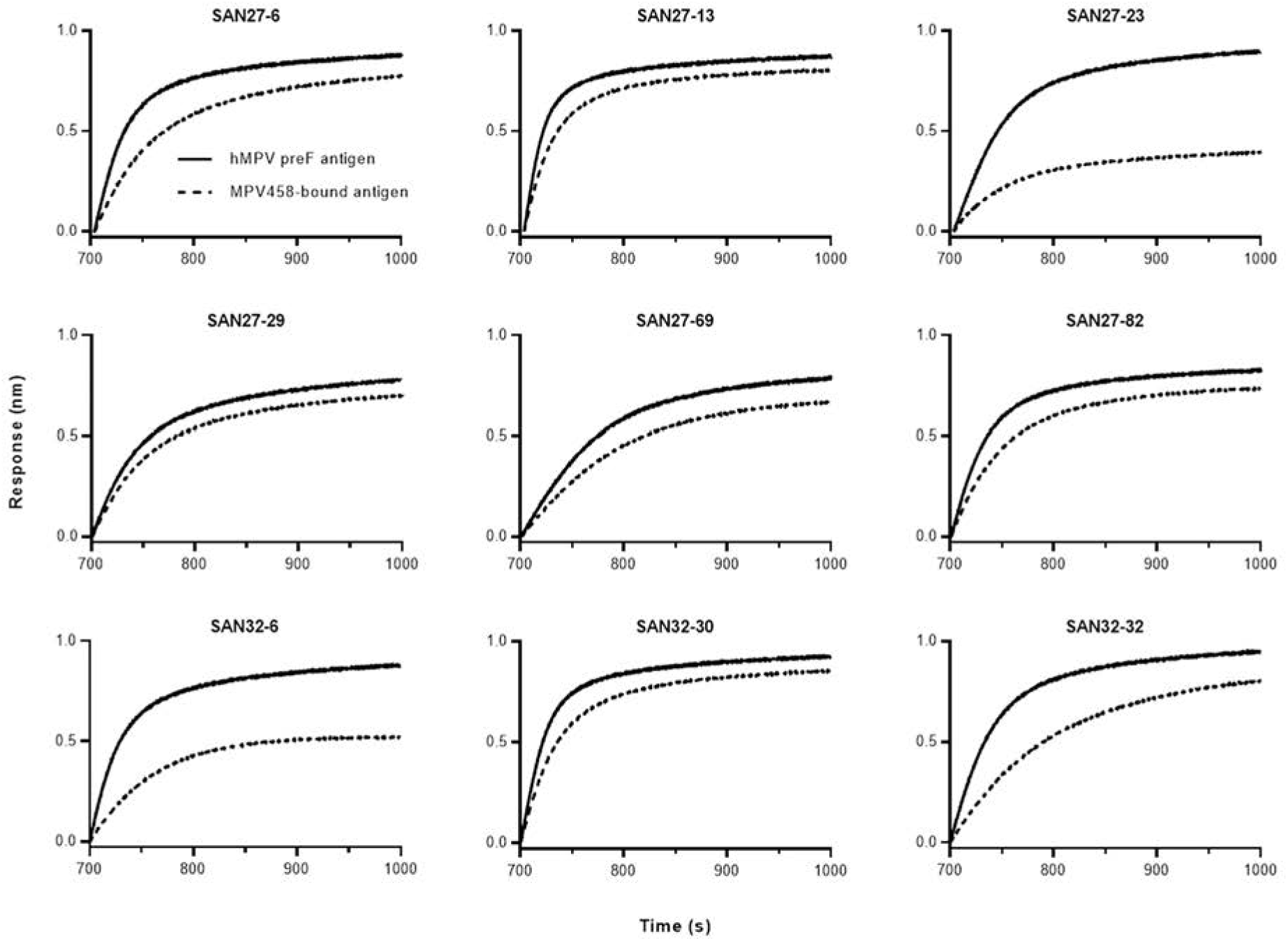
Bin A mAbs partially compete with trimer-interface Fab MPV458. Biolayer interferometry assay sensorgrams showing indicated antibody’s binding association profile to 115-BV-DS (solid line) and to 115-BV-DS after it has been bound by MPV458 Fab (dashed line).

**Supplementary Figure 4.**
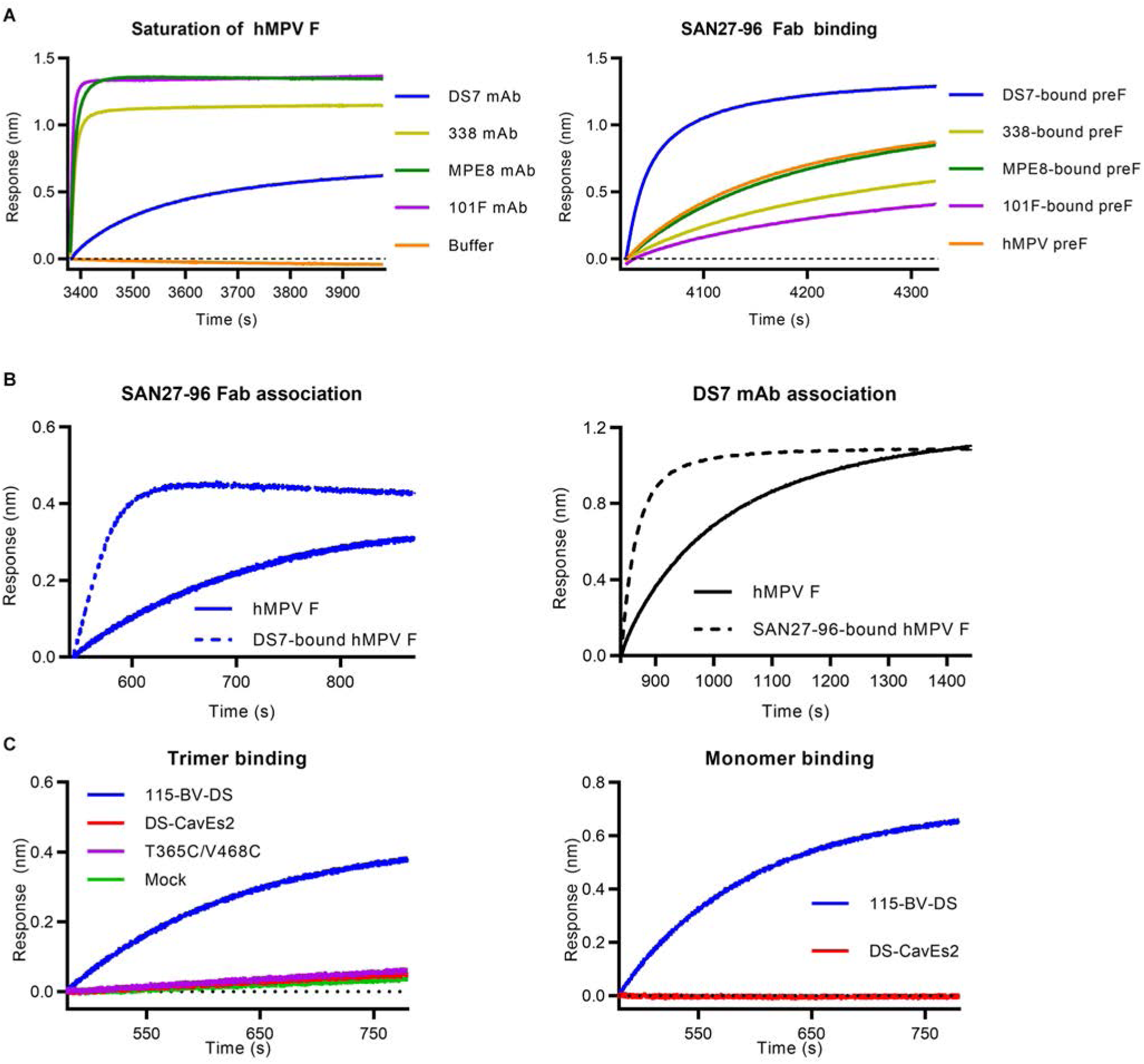
Representative Bin D antibody SAN27-96 competition profile. **A.** Biolayer interferometry (BLI) sensorgrams showing hMPV F 115-BV-DS being saturated with primary antibody (left) and the secondary SAN27-96 Fab binding signal to saturated 115-BV-DS (right). **B**. BLI curves showing the enhanced antibody association seen for Bin D when hMPV F is first saturated with DS7 mAb (left). The same enhanced association is also seen for DS7 mAb when hMPV F is first saturated with representative Bin D Fab SAN27-96(right). **C.** BLI curves of SAN27-96 binding to a panel of hMPV F trimer (left) and monomer (right) constructs.

**Supplementary Figure 5.**
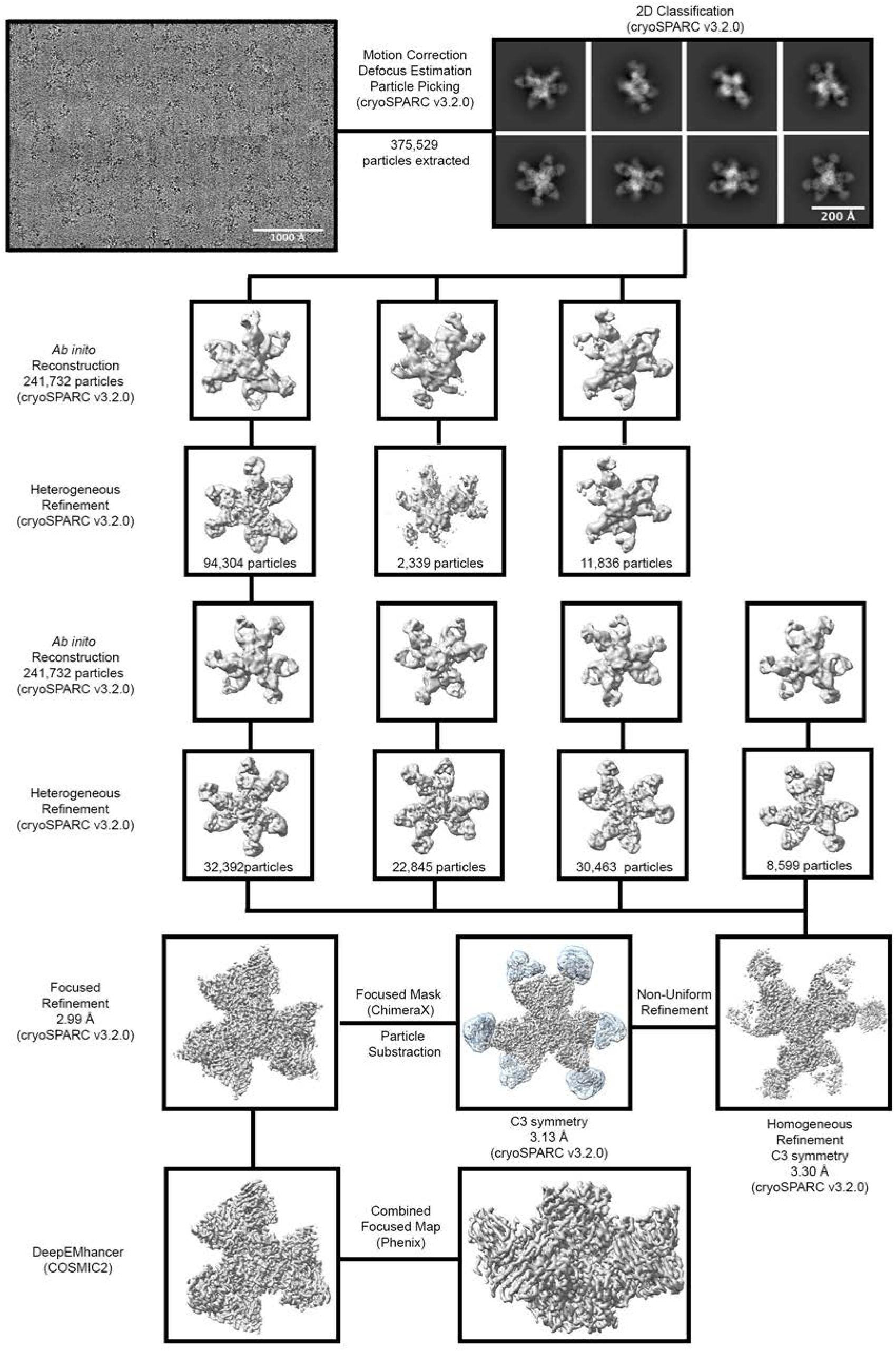
SAN27-14 cryo-EM processing workflow. Each step, from representative micrograph to combined focus map, of the cryo-EM data processing workflow is shown. Computational programs and algorithms used are labeled for each step. The mask used for particle subtraction is colored as a transparent cyan.

**Supplementary Figure 6.**
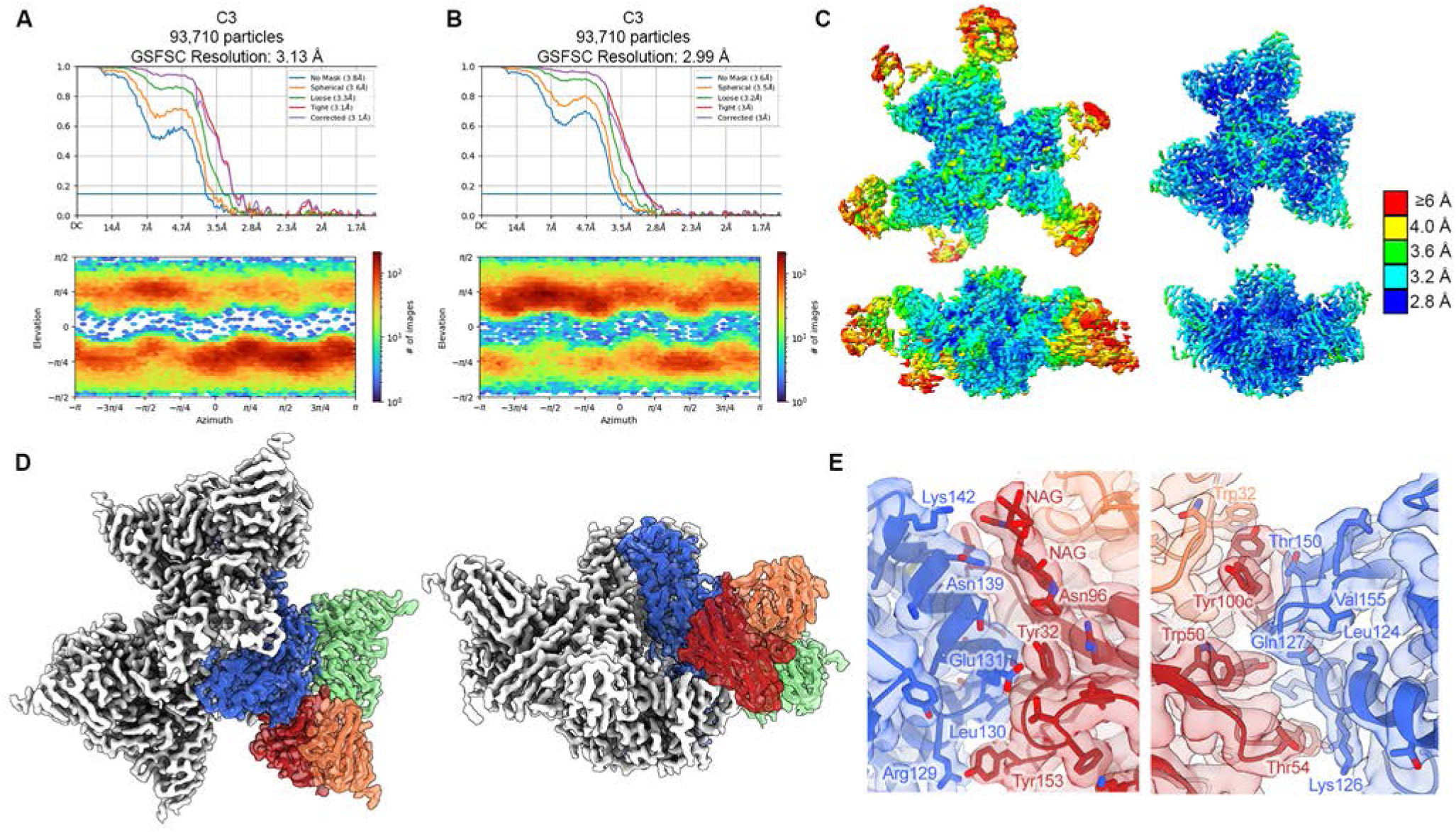
SAN27-14 cryo-EM validation. **A.** (top) The FSC curves for the non-uniform refinement 3D reconstruction. The horizontal blue line corresponds to an FSC value of 0.143. (bottom) The viewing distribution plot calculated in cryoSPARC is shown. **B.** The FSC curves and viewing distribution plot for the focused refinement 3D reconstruction is shown. (top) The FSC curves for the non-uniform refinement 3D reconstruction. The horizontal blue line corresponds to an FSC value of 0.143. (bottom) The viewing distribution plot calculated in cryoSPARC is shown. **C.** Cryo-EM maps of non-uniform refinement (left) and focused refinement(right) colored by local resolution. Cryo-EM maps are shown as top (top) and side (bottom) views. **D.** Cryo-EM map of DS-CavEs2 bound by MPE8 Fab and SAN27-14 is shown in a top view (left) and side view(right). A single protomer is shown as a transparent colored surface with a docked ribbon model (DS-CavEs2: blue; SAN27-14: red, orange; MPE8: green). **E.** The binding interface for SAN27-14 with DS-CavEs2. Cryo-EM map is shown as a transparent surface with the docked model shown as a ribbon. Colored the same as in **D.**

**Supplementary Figure 7.**
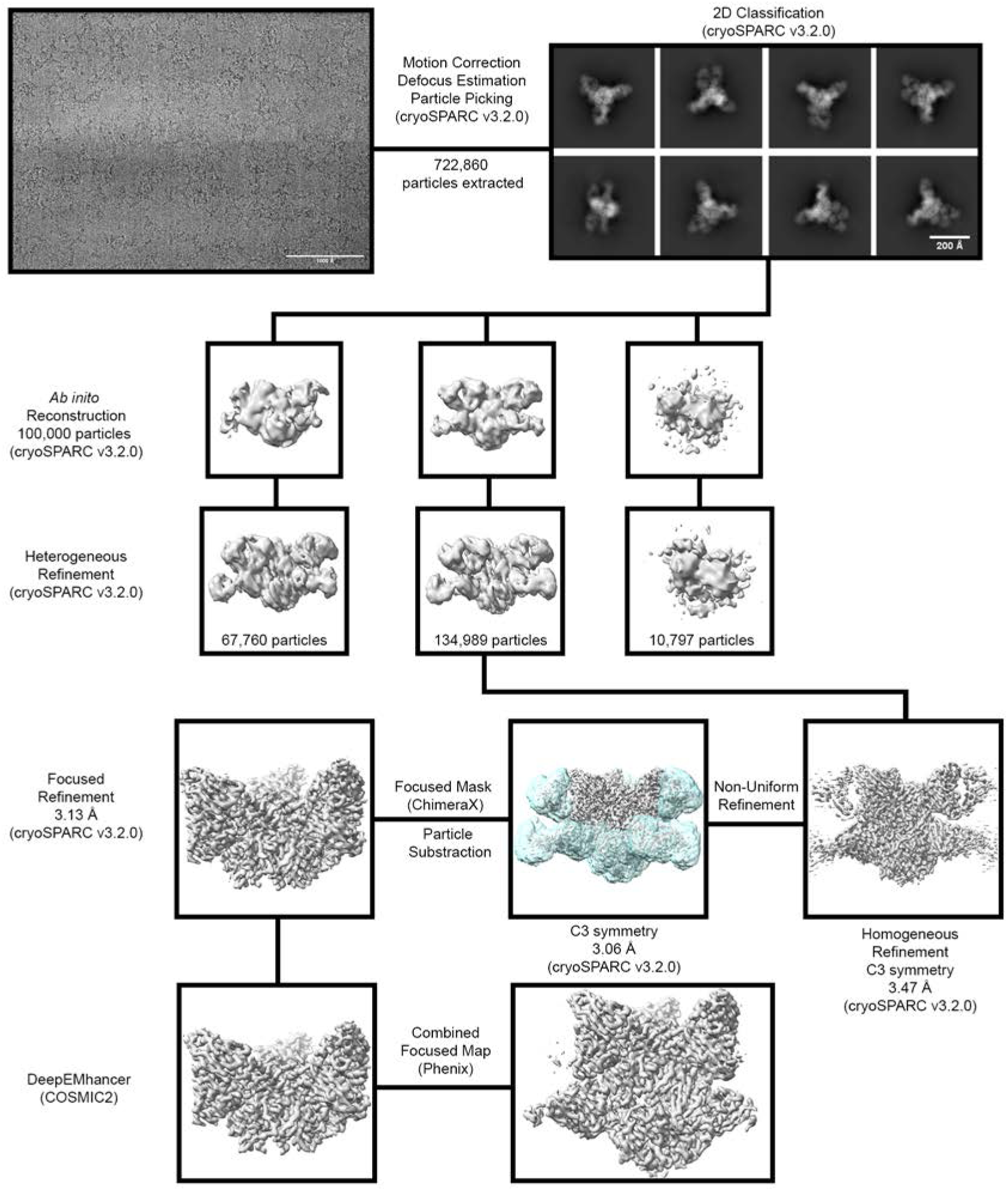
SAN32-2 cryo-EM processing workflow. Each step, from representative micrograph to combined focus map, of the cryo-EM data processing workflow is shown. Computational programs and algorithms used are labeled for each step. The mask used for particle subtraction is colored as a transparent cyan.

**Supplementary Figure 8.**
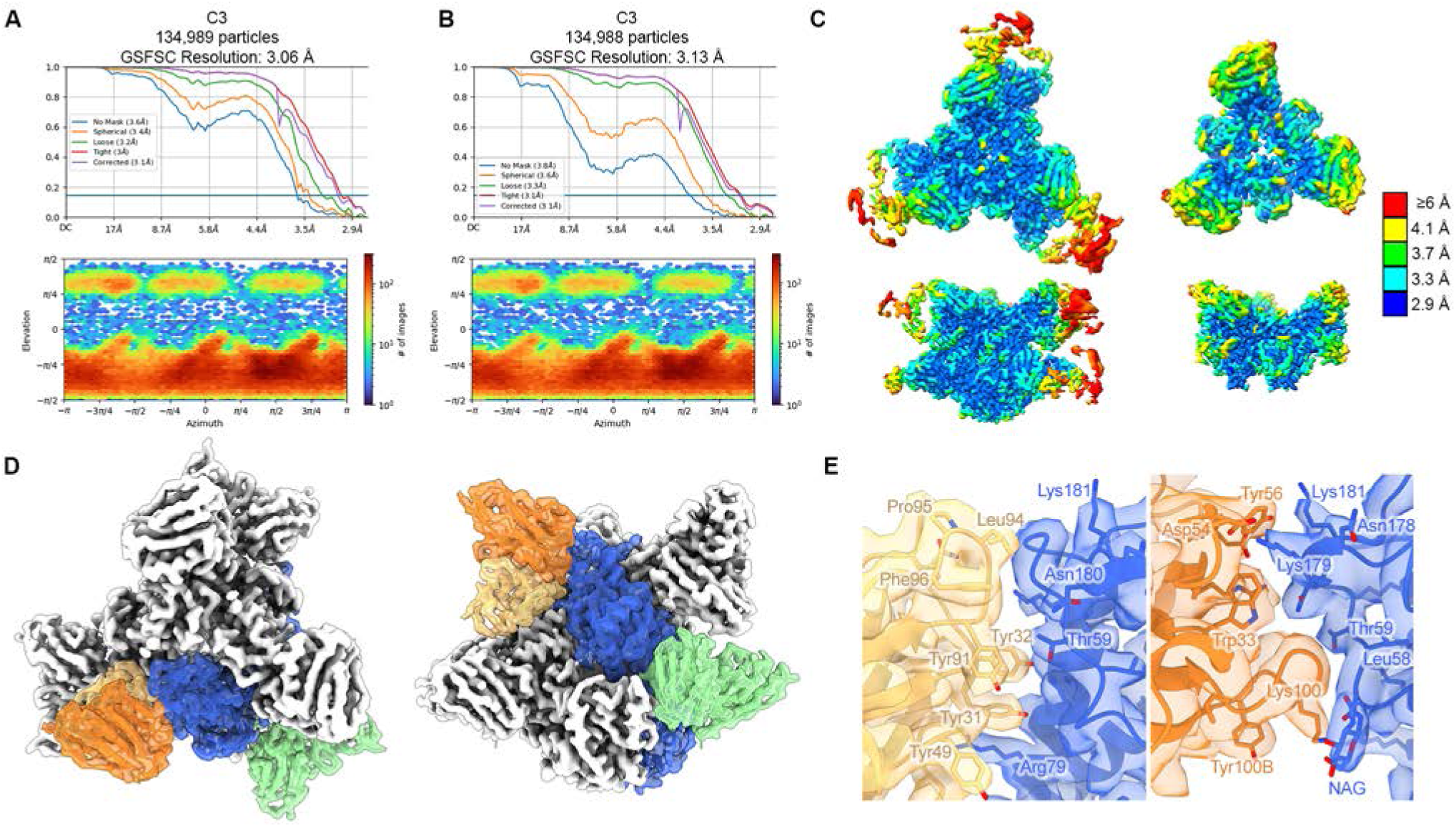
SAN32-2 cryo-EM Validation. **A.** (top) The FSC curves for the non-uniform refinement 3D reconstruction. The horizontal blue line corresponds to an FSC value of 0.143. (bottom) The viewing distribution plot calculated in cryoSPARC is shown. **B.** The FSC curves and viewing distribution plot for the focused refinement 3D reconstruction is shown. (top) The FSC curves for the non-uniform refinement 3D reconstruction. The horizontal blue line corresponds to an FSC value of 0.143. (bottom) The viewing distribution plot calculated in cryoSPARC is shown. **C.** Cryo-EM maps of non-uniform refinement (left) and focused refinement(right) colored by local resolution. Cryo-EM maps are shown as top (top) and side (bottom) views. **D.** Cryo-EM map of DS-CavEs2 bound by MPE8 Fab and SAN32-2 is shown in a top view (left) and side view(right). A single protomer is shown as a transparent colored surface with a docked ribbon model (DS-CavEs2: blue; SAN32-2: orange, light orange; MPE8: green). **E.** The binding interface for SAN32-2 light chain (left) and heavy chain (right) with DS-CavEs2. Cryo-EM map is shown as a transparent surface with the docked model shown as a ribbon. Colored the same as in **D.**

**Supplemental Table 1.** hMPV Prefusion F ELISA titer in plasma from elderly donors

| Donor | Gender | Donation | Date of Donation | Age at Donation | hMPV Prefusion F ELISA Titer |
| --- | --- | --- | --- | --- | --- |
| 1 | M | 1.1 | 7/7/2014 | 52 | 8000 |
| 2 | M | 2.1 | 8/5/2013 | 65 | 8000 |
|  |  | 2.2 | 6/12/2014 | 66 | 16000 |
|  |  | 2.3 | 5/22/2015 | 67 | 64000 |
|  |  | 2.4 | 5/17/2016 | 68 | 32000 |
|  |  | 2.5 | 3/28/2017 | 68 | 16000 |
| 3 | M | 3.1 | 2/15/2018 | 68 | 16000 |
|  |  | 3.2 | 4/18/2018 | 68 | 4000 |
|  |  | 3.3 | 5/13/2019 | 69 | 4000 |
| 4 | M | 4.1 | 7/200/17 | 59 | 4000 |
|  |  | 4.2 | 1/17/2018 | 59 | 64000 |
|  |  | 4.3 | 4/11/2019 | 61 | 32000 |
| 5 | F | 5.1 | 4/27/2015 | 56 | 16000 |
|  |  | 5.2 | 5/4/2016 | 57 | 16000 |
|  |  | 5.3 | 10/5/2017 | 59 | 32000 |
| 6 | F | 6.1 | 6/25/2013 | 66 | 16000 |
|  |  | 6.2 | 9/9/2013 | 66 | 32000 |
|  |  | 6.3 | 9/20/2018 | 71 | 32000 |
| 7 | M | 7.1 | 10/11/2017 | 63 | 16000 |
| 8 | M | 8.1 | 8/14/2013 | 69 | 32000 |
|  |  | 8.2 | 4/2/2014 | 70 | 16000 |
|  |  | 8.3 | 8/25/2014 | 70 | 16000 |
|  |  | 8.4 | 4/24/2015 | 71 | 8000 |
| 9 | F | 9.1 | 1/15/2014 | 74 | 32000 |
|  |  | 9.2 | 3/16/2015 | 76 | 32000 |
| 10 | M | 10.1 | 5/8/2019 | 73 | 16000 |
| 11 | F | 11.1 | 12/18/2014 | 68 | 32000 |
|  |  | 11.2 | 3/4/2019 | 72 | 64000 |
|  |  | 11.3 | 5/6/2019 | 72 | 16000 |
| 12 | M | 12.1 | 5/16/2019 | 70 | 8000 |
|  |  | 12.2 | 10/28/2019 | 70 | 16000 |

**Supplemental Table 2.**
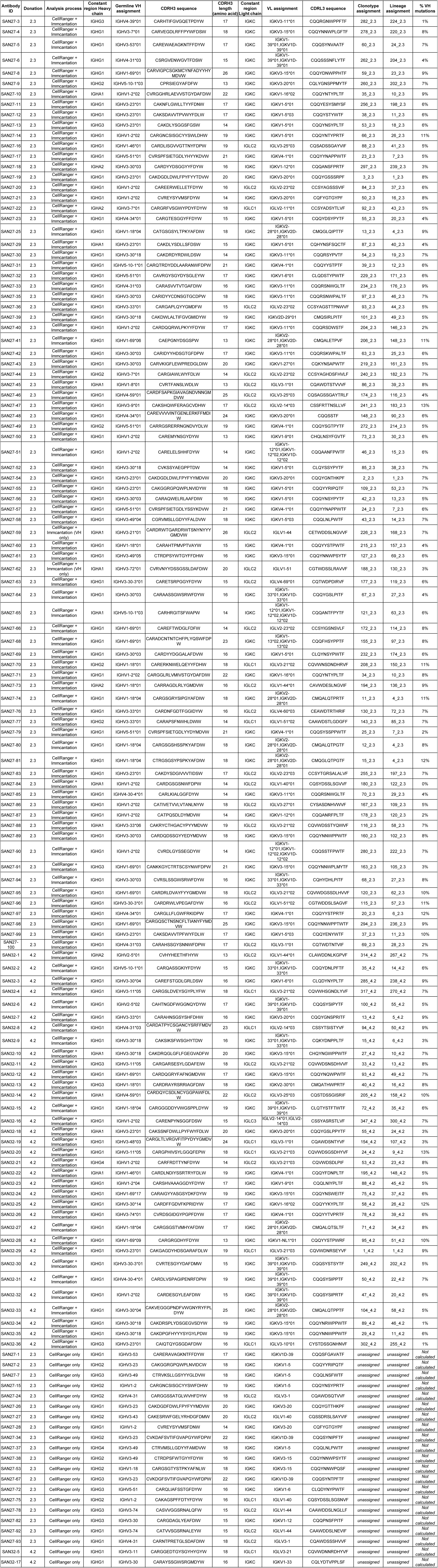
Sequence features, germline assignments, and clustering analysis of expressed mAbs.

**Supplemental Table 3.** hMPV binding and neutralization properties of expressed mAbs.

| Antibody ID | Donation | A2 Neut (µg/mL) | hMPV A1 PreF ELISA (ng/mL) | hMPV A1 PostF ELISA (ng/mL)2 | hMPV B2 PreF ELISA (ng/mL) |
| --- | --- | --- | --- | --- | --- |
| SAN27-1 | 2.3 | 0.44 | 7.1 | 111 | 1.4 |
| SAN27-2 | 2.3 | 0.82 | 2.4 | 4 | 0.5 |
| SAN27-3 | 2.3 | >12.5 | >1000 | >1000 | >1000 |
| SAN27-4 | 2.3 | >12.5 | 111.1 | ND | ND |
| SAN27-5 | 2.3 | 0.54 | 4.1 | 12 | 0.2 |
| SAN27-6 | 2.3 | 1.85 | 2.4 | 1 | 0.5 |
| SAN27-7 | 2.3 | 0.06 | 2.4 | 1 | 0.2 |
| SAN27-8 | 2.3 | 0.98 | 2.4 | 12 | 0.5 |
| SAN27-9 | 2.3 | >12.5 | >1000 | >1000 | >1000 |
| SAN27-10 | 2.3 | 0.07 | 0.9 | 333 | 0.2 |
| SAN27-11 | 2.3 | 0.32 | 0.5 | 12 | 0.2 |
| SAN27-12 | 2.3 | 0.27 | 4.1 | 333 | 0.2 |
| SAN27-13 | 2.3 | 1.02 | 0.9 | 4 | 0.2 |
| SAN27-14 | 2.3 | 0.28 | 2.4 | 37 | 0.2 |
| SAN27-15 | 2.3 | 0.32 | 2.4 | 37 | 0.2 |
| SAN27-16 | 2.3 | >12.5 | >1000 | >1000 | >1000 |
| SAN27-17 | 2.3 | 1.62 | 12.3 | 333 | 0.4 |
| SAN27-18 | 2.3 | >12.5 | >1000 | >1000 | >1000 |
| SAN27-19 | 2.3 | 0.30 | 4.1 | 111 | 0.2 |
| SAN27-20 | 2.3 | 3.74 | 4.1 | 111 | 0.2 |
| SAN27-21 | 2.3 | 3.38 | 2.4 | 37 | 0.5 |
| SAN27-22 | 2.3 | >12.5 | >1000 | >1000 | >1000 |
| SAN27-23 | 2.3 | 0.99 | 2.4 | 4 | 0.5 |
| SAN27-24 | 2.3 | 0.64 | 2.4 | 12 | 0.5 |
| SAN27-25 | 2.3 | >12.5 | 577.4 | >1000 | 333.3 |
| SAN27-26 | 2.3 | 0.47 | 2.4 | 111 | 0.2 |
| SAN27-27 | 2.3 | 0.11 | 12.3 | >1000 | 0.4 |
| SAN27-28 | 2.3 | 1.63 | 7.1 | 37 | 0.2 |
| SAN27-29 | 2.3 | 1.08 | 0.9 | 12 | 0.2 |
| SAN27-30 | 2.3 | 0.95 | 2.4 | 1000 | 0.2 |
| SAN27-31 | 2.3 | >12.5 | 4.1 | >1000 | 4.1 |
| SAN27-32 | 2.3 | >12.5 | >1000 | >1000 | >1000 |
| SAN27-33 | 2.3 | 0.09 | 2.4 | 1 | 0.2 |
| SAN27-34 | 2.3 | 1.13 | 2.4 | 4 | 0.2 |
| SAN27-35 | 2.3 | 0.28 | 1.4 | 4 | 0.2 |
| SAN27-36 | 2.3 | 1.57 | 7.1 | 111 | 0.5 |
| SAN27-37 | 2.3 | 0.07 | 2.4 | 4 | 0.2 |
| SAN27-38 | 2.3 | 0.84 | 64.2 | 37 | 0.2 |
| SAN27-39 | 2.3 | 4.14 | 64.2 | >1000 | 2.4 |
| SAN27-40 | 2.3 | 1.19 | 1.6 | 37 | 0.2 |
| SAN27-41 | 2.3 | 0.09 | 0.9 | 12 | 0.2 |
| SAN27-42 | 2.3 | 0.20 | 12.3 | 4 | 0.2 |
| SAN27-43 | 2.3 | >12.5 | >1000 | 333 | >1000 |
| SAN27-44 | 2.3 | >12.5 | >1000 | >1000 | >1000 |
| SAN27-45 | 2.3 | 1.68 | 37.0 | >1000 | 0.2 |
| SAN27-46 | 2.3 | >12.5 | >1000 | >1000 | >1000 |
| SAN27-47 | 2.3 | 0.03 | 4.1 | 1 | 0.2 |
| SAN27-48 | 2.3 | >12.5 | 4.1 | ND | ND |
| SAN27-49 | 2.3 | >12.5 | >1000 | >1000 | >1000 |
| SAN27-50 | 2.3 | 2.61 | 111.1 | 111 | 2.4 |
| SAN27-51 | 2.3 | 3.46 | 4.1 | 12 | 2.4 |
| SAN27-52 | 2.3 | 1.02 | 0.5 | 4 | 0.5 |
| SAN27-53 | 2.3 | >12.5 | >1000 | >1000 | >1000 |
| SAN27-54 | 2.3 | 0.51 | 0.2 | 37 | 0.5 |
| SAN27-55 | 2.3 | 0.93 | 7.1 | 12 | 0.2 |
| SAN27-56 | 2.3 | 0.30 | 0.2 | 12 | 1.4 |
| SAN27-57 | 2.3 | 1.10 | 0.9 | 111 | 1.4 |
| SAN27-58 | 2.3 | 0.07 | 0.2 | 4 | 0.2 |
| SAN27-59 | 2.3 | >12.5 | 37.0 | 12 | 12.3 |
| SAN27-60 | 2.3 | >12.5 | 64.2 | >1000 | >1000 |
| SAN27-61 | 2.3 | 0.82 | 2.4 | 4 | 0.5 |
| SAN27-62 | 2.3 | >12.5 | >1000 | >1000 | >1000 |
| SAN27-63 | 2.3 | 0.27 | 0.5 | 1 | 0.2 |
| SAN27-64 | 2.3 | 0.05 | 12.3 | <0.4 | 0.5 |
| SAN27-65 | 2.3 | 0.22 | 0.9 | 12 | 0.2 |
| SAN27-66 | 2.3 | 3.15 | 0.9 | 37 | 0.5 |
| SAN27-67 | 2.3 | 1.03 | 0.5 | 1 | 0.2 |
| SAN27-68 | 2.3 | 3.19 | 4.1 | 37 | 1.4 |
| SAN27-69 | 2.3 | 1.29 | 0.9 | 4 | 0.2 |
| SAN27-70 | 2.3 | >12.5 | 333.3 | >1000 | 37.0 |
| SAN27-71 | 2.3 | 0.20 | 4.1 | 12 | 0.5 |
| SAN27-72 | 2.3 | >12.5 | >1000 | >1000 | >1000 |
| SAN27-73 | 2.3 | >12.5 | >1000 | >1000 | >1000 |
| SAN27-74 | 2.3 | >12.5 | 0.9 | 12 | 0.2 |
| SAN27-75 | 2.3 | 1.94 | 21.4 | 37 | 4.1 |
| SAN27-76 | 2.3 | 1.97 | 0.9 | 4 | 1.4 |
| SAN27-77 | 2.3 | >12.5 | >1000 | >1000 | >1000 |
| SAN27-78 | 2.3 | >12.5 | >1000 | >1000 | >1000 |
| SAN27-79 | 2.3 | 1.24 | 7.1 | 111 | 1.4 |
| SAN27-80 | 2.3 | 0.59 | 192.5 | >1000 | 37.0 |
| SAN27-81 | 2.3 | >12.5 | 333.3 | >1000 | 37.0 |
| SAN27-82 | 2.3 | 1.69 | 0.5 | <0.4 | 0.5 |
| SAN27-83 | 2.3 | 6.97 | 7.1 | 37 | 2.4 |
| SAN27-84 | 2.3 | >12.5 | >1000 | >1000 | >1000 |
| SAN27-85 | 2.3 | 0.05 | 0.9 | 4 | 0.5 |
| SAN27-86 | 2.3 | >12.5 | 0.9 | <0.4 | 1.4 |
| SAN27-87 | 2.3 | 1.37 | 0.5 | 12 | 0.5 |
| SAN27-88 | 2.3 | >12.5 | >1000 | >1000 | >1000 |
| SAN27-89 | 2.3 | >12.5 | >1000 | >1000 | >1000 |
| SAN27-90 | 2.3 | 3.74 | 0.5 | 12 | 1.4 |
| SAN27-91 | 2.3 | >12.5 | 37.0 | ND | ND |
| SAN27-92 | 2.3 | >12.5 | 333.3 | ND | ND |
| SAN27-93 | 2.3 | 0.99 | 0.2 | <0.4 | 0.2 |
| SAN27-94 | 2.3 | 0.09 | 0.2 | <0.4 | 0.2 |
| SAN27-95 | 2.3 | >12.5 | 0.2 | <0.4 | 0.2 |
| SAN27-96 | 2.3 | 0.98 | 0.2 | <0.4 | 0.2 |
| SAN27-97 | 2.3 | 0.07 | 14.1 | <0.4 | 0.2 |
| SAN27-98 | 2.3 | 1.21 | 0.2 | ND | ND |
| SAN27-99 | 2.3 | 0.27 | 0.2 | ND | 0.2 |
| SAN27-100 | 2.3 | 1.46 | 0.2 | ND | 0.2 |
| SAN32-1 | 4.2 | >12.5 | 577.0 | >1000 | >1000 |
| SAN32-2 | 4.2 | 0.08 | 2.0 | 1 | <0.4 |
| SAN32-3 | 4.2 | >12.5 | >1000 | >1000 | >1000 |
| SAN32-4 | 4.2 | 0.22 | 2.0 | 7 | <0.4 |
| SAN32-5 | 4.2 | 0.26 | 2.0 | 21 | <0.4 |
| SAN32-6 | 4.2 | 5.25 | 1.0 | 4 | <0.4 |
| SAN32-7 | 4.2 | 0.14 | 2.0 | 21 | <0.4 |
| SAN32-8 | 4.2 | 0.71 | 4.0 | 2 | <0.4 |
| SAN32-9 | 4.2 | 0.06 | 4.0 | 1 | <0.4 |
| SAN32-10 | 4.2 | >12.5 | 2.0 | 12 | 1.0 |
| SAN32-11 | 4.2 | 2.01 | 2.0 | 64 | <0.4 |
| SAN32-12 | 4.2 | 0.24 | 2.0 | 1 | <0.4 |
| SAN32-13 | 4.2 | >12.5 | >1000 | >1000 | >1000 |
| SAN32-14 | 4.2 | >12.5 | >1000 | >1000 | >1000 |
| SAN32-15 | 4.2 | 1.47 | 4.0 | 2 | <0.4 |
| SAN32-16 | 4.2 | 4.77 | 7.0 | 12 | 1.0 |
| SAN32-17 | 4.2 | 0.23 | 4.0 | 12 | 1.0 |
| SAN32-18 | 4.2 | 0.76 | 4.0 | 21 | <0.4 |
| SAN32-19 | 4.2 | 1.38 | 4.0 | 12 | <0.4 |
| SAN32-20 | 4.2 | 0.25 | 2.0 | 4 | <0.4 |
| SAN32-21 | 4.2 | >12.5 | >1000 | >1000 | >1000 |
| SAN32-22 | 4.2 | >12.5 | >1000 | >1000 | >1000 |
| SAN32-23 | 4.2 | 2.63 | 7.0 | 64 | 1.0 |
| SAN32-24 | 4.2 | 0.36 | 2.0 | 4 | <0.4 |
| SAN32-25 | 4.2 | 0.94 | 7.00 | <0.4 | <0.4 |
| SAN32-26 | 4.2 | 9.87 | >1000 | >1000 | >1000 |
| SAN32-27 | 4.2 | >12.5 | >1000 | >1000 | >1000 |
| SAN32-28 | 4.2 | >12.5 | 12.00 | 7.00 | 3.00 |
| SAN32-29 | 4.2 | 0.07 | 1.00 | <0.4 | <0.4 |
| SAN32-30 | 4.2 | 1.23 | 4.00 | 1.00 | <0.4 |
| SAN32-31 | 4.2 | 0.88 | 2.00 | 4.00 | <0.4 |
| SAN32-32 | 4.2 | 1.35 | 1.00 | <0.4 | <0.4 |
| SAN32-33 | 4.2 | 3.35 | 1.00 | <0.4 | <0.4 |
| SAN32-34 | 4.2 | 2.55 | 1.00 | <0.4 | <0.4 |
| SAN32-35 | 4.2 | 0.60 | <0.4 | <0.4 | <0.4 |
| SAN32-36 | 4.2 | >12.5 | >1000 | >1000 | >1000 |

**Supplemental Table 4.**
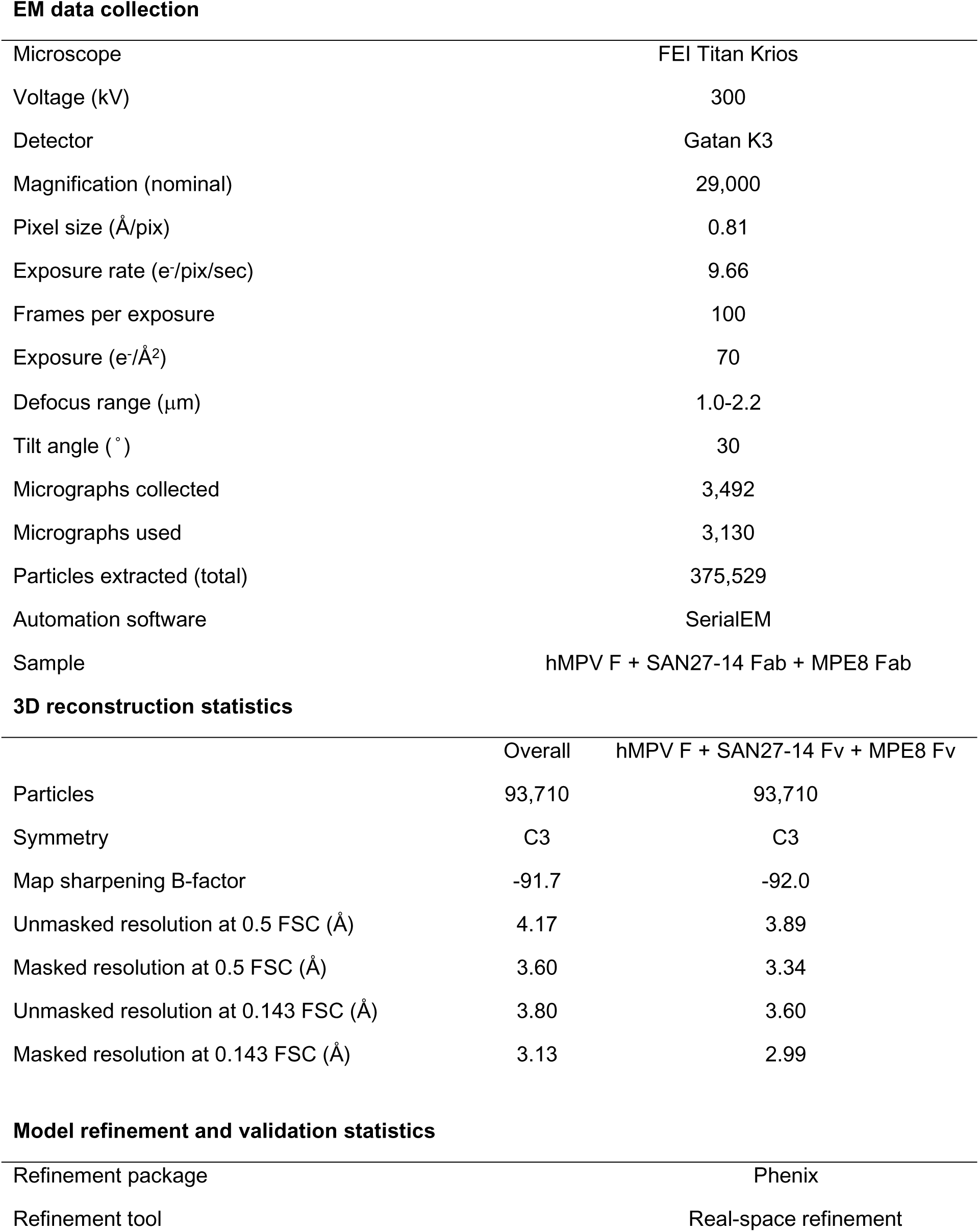
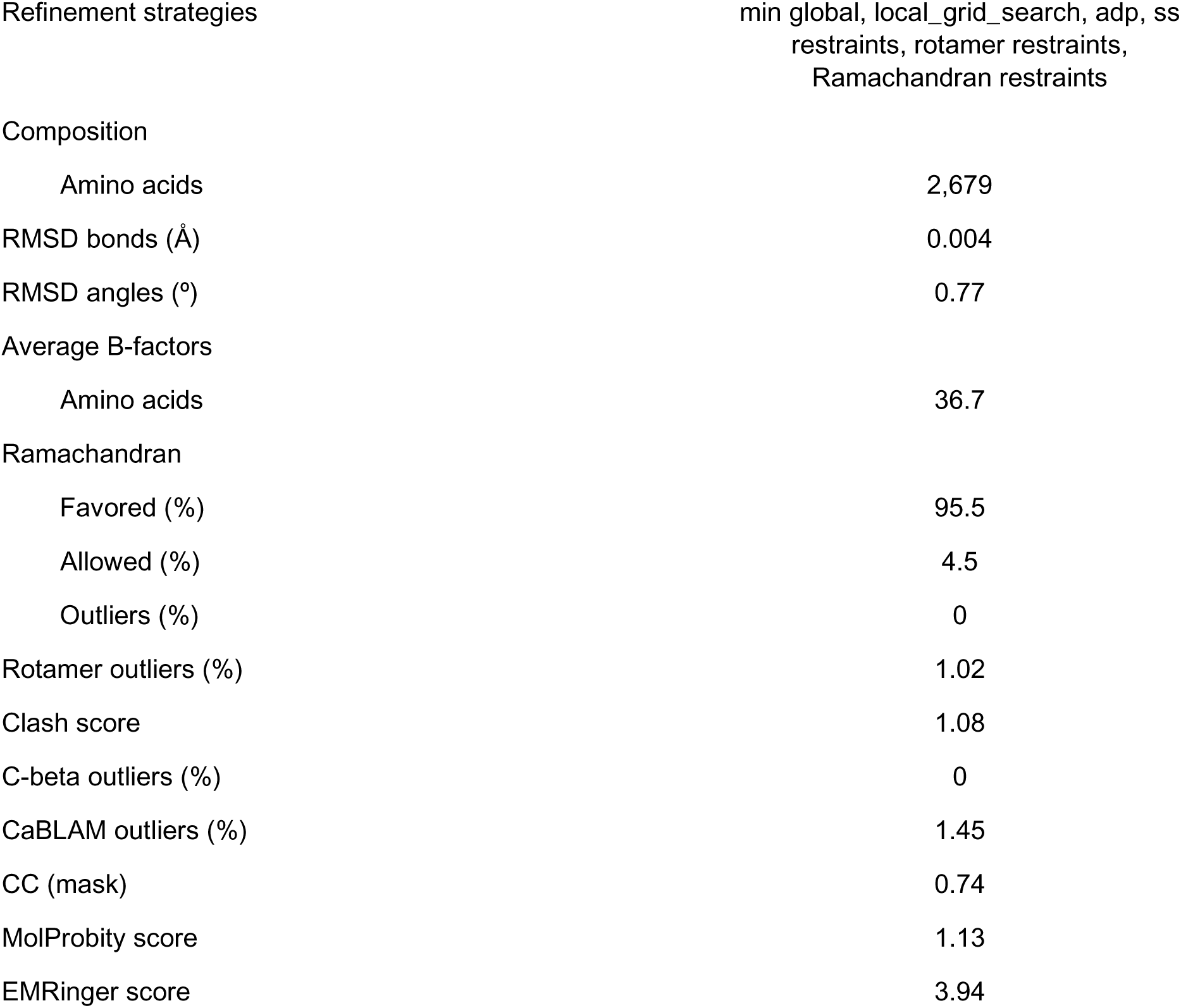
EM data collection for SAN27-14

|  |  |
| --- | --- |
| Microscope | FEI Titan Krios |
| Voltage (kV) | 300 |
| Detector | Gatan K3 |
| Magnification (nominal) | 29,000 |
| Pixel size (Å/pix) | 0.81 |
| Exposure rate (e<sup>-</sup>/pix/sec) | 9.66 |
| Frames per exposure | 100 |
| Exposure (e<sup>-</sup>/Å<sup>2</sup>) | 70 |
| Defocus range (μm) | 1.0-2.2 |
| Tilt angle (°) | 30 |
| Micrographs collected | 3,492 |
| Micrographs used | 3,130 |
| Particles extracted (total) | 375,529 |
| Automation software | SerialEM |
| Sample | hMPV F + SAN27-14 Fab + MPE8 Fab |

**3D reconstruction statistics**
|  | Overall | hMPV F + SAN27-14 Fv + MPE8 Fv |
| --- | --- | --- |
| Particles | 93,710 | 93,710 |
| Symmetry | C3 | C3 |
| Map sharpening B-factor | -91.7 | -92.0 |
| Unmasked resolution at 0.5 FSC (Å) | 4.17 | 3.89 |
| Masked resolution at 0.5 FSC (Å) | 3.60 | 3.34 |
| Unmasked resolution at 0.143 FSC (Å) | 3.80 | 3.60 |
| Masked resolution at 0.143 FSC (Å) | 3.13 | 2.99 |
| **Model refinement and validation statistics** |  |
| Refinement package | Phenix |
| Refinement tool | Real-space refinement |
| Composition |  |
| Amino acids | 2,679 |
| RMSD bonds (Å) | 0.004 |
| RMSD angles (°) | 0.77 |
| Average B-factors |  |
| Amino acids | 36.7 |
| Ramachandran |  |
| Favored (%) | 95.5 |
| Allowed (%) | 4.5 |
| Outliers (%) | 0 |
| Rotamer outliers (%) | 1.02 |
| Clash score | 1.08 |
| C-beta outliers (%) | 0 |
| CaBLAM outliers (%) | 1.45 |
| CC (mask) | 0.74 |
| MolProbity score | 1.13 |
| EMRinger score | 3.94 |

**Supplemental Table 5.**
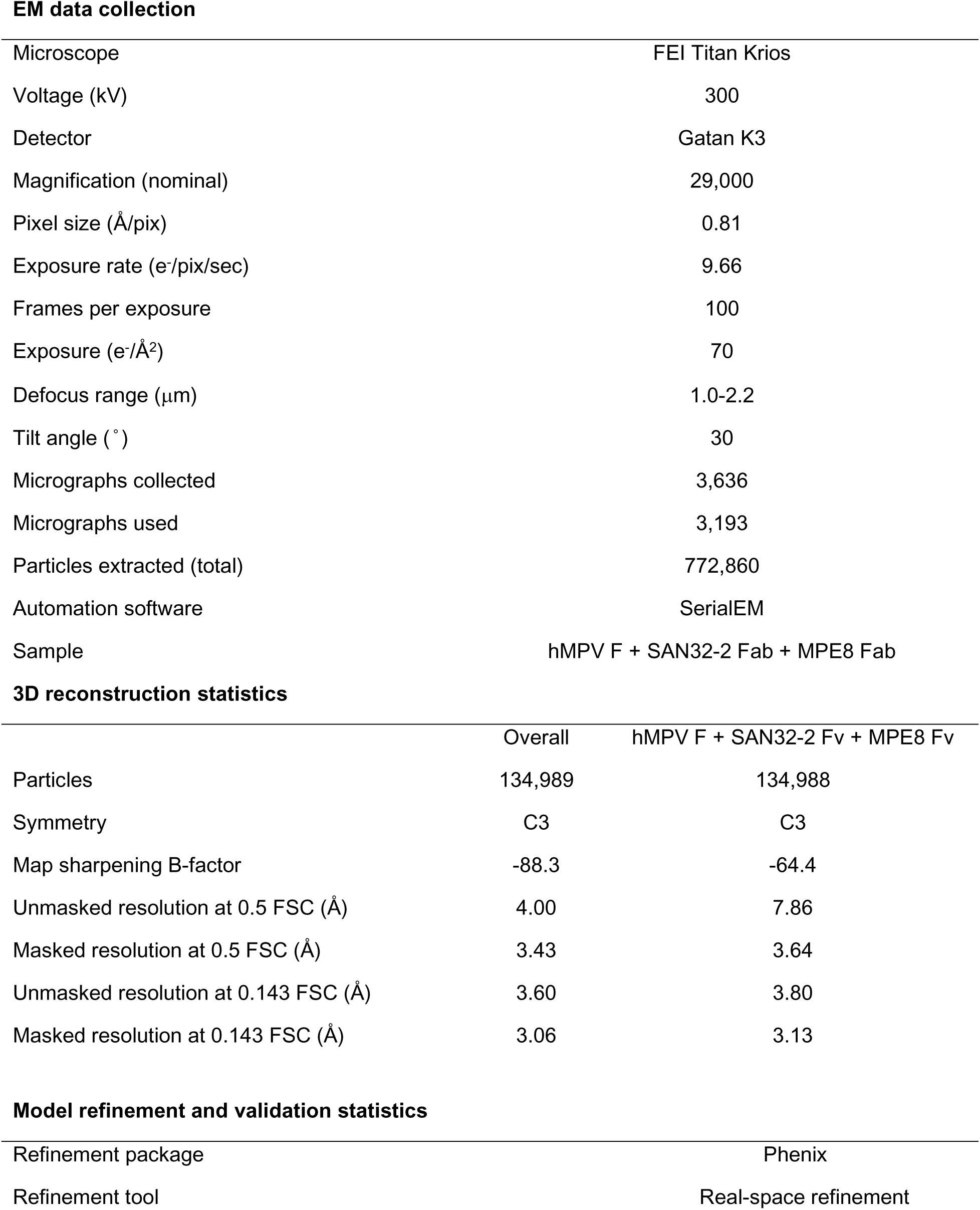
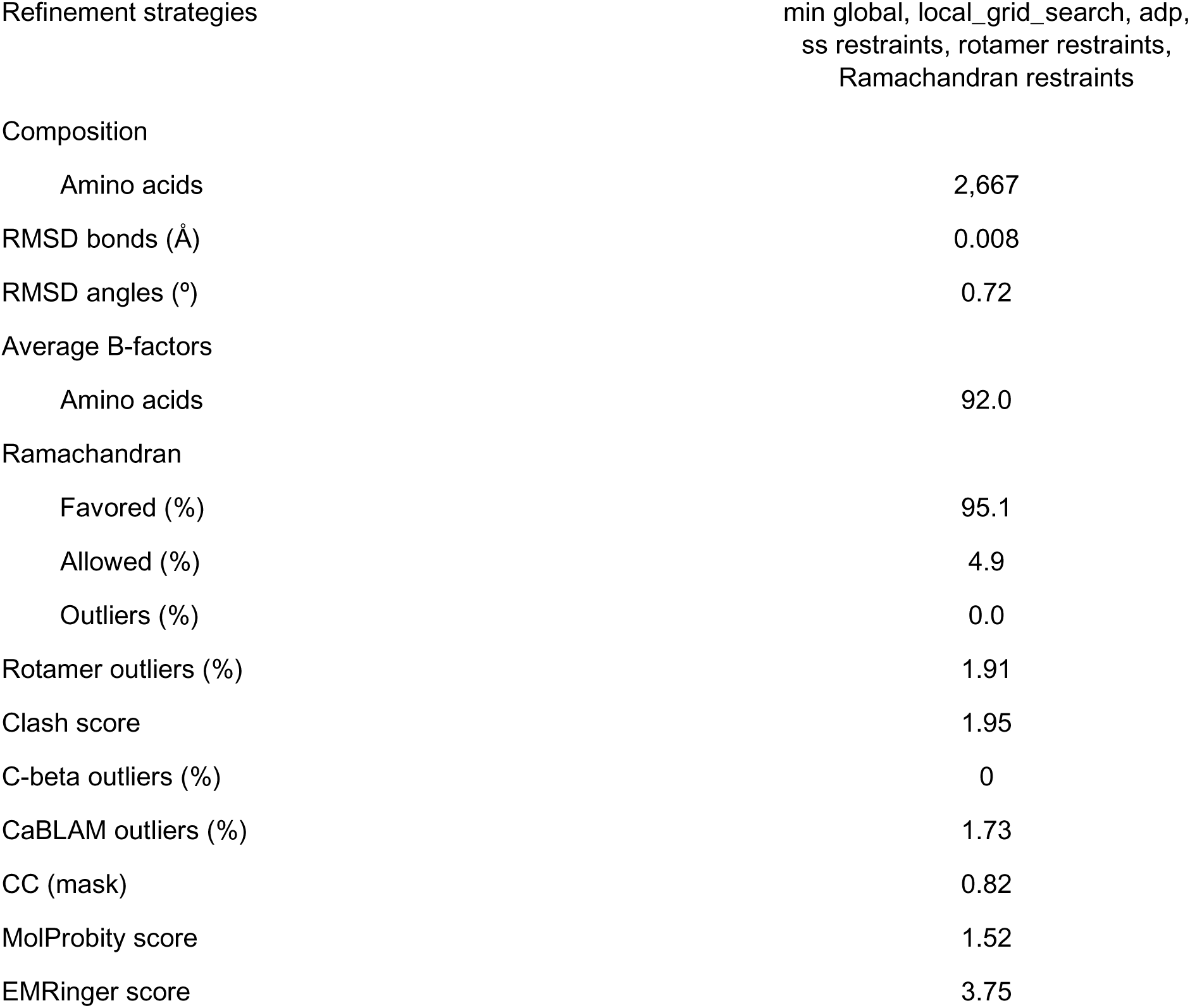
EM data collection for SAN32-2

|  |  |
| --- | --- |
| Microscope | FEI Titan Krios |
| Voltage (kV) | 300 |
| Detector | Gatan K3 |
| Magnification (nominal) | 29,000 |
| Pixel size (Å/pix) | 0.81 |
| Exposure rate (e<sup>-</sup>/pix/sec) | 9.66 |
| Frames per exposure | 100 |
| Exposure (e<sup>-</sup>/Å<sup>2</sup>) | 70 |
| Defocus range (μm) | 1.0-2.2 |
| Tilt angle (°) | 30 |
| Micrographs collected | 3,636 |
| Micrographs used | 3,193 |
| Particles extracted (total) | 772,860 |
| Automation software | SerialEM |
| Sample | hMPV F + SAN32-2 Fab + MPE8 Fab |

**3D reconstruction statistics**
|  | Overall | hMPV F + SAN32-2 Fv + MPE8 Fv |
| --- | --- | --- |
| Particles | 134,989 | 134,988 |
| Symmetry | C3 | C3 |
| Map sharpening B-factor | -88.3 | -64.4 |
| Unmasked resolution at 0.5 FSC (Å) | 4.00 | 7.86 |
| Masked resolution at 0.5 FSC (Å) | 3.43 | 3.64 |
| Unmasked resolution at 0.143 FSC (Å) | 3.60 | 3.80 |
| Masked resolution at 0.143 FSC (Å) | 3.06 | 3.13 |
| **Model refinement and validation statistics** |  |
| Refinement package | Phenix |
| Refinement tool | Real-space refinement |
| Composition |  |
| Amino acids | 2,667 |
| RMSD bonds (Å) | 0.008 |
| RMSD angles (°) | 0.72 |
| Average B-factors |  |
| Amino acids | 92.0 |
| Ramachandran |  |
| Favored (%) | 95.1 |
| Allowed (%) | 4.9 |
| Outliers (%) | 0.0 |
| Rotamer outliers (%) | 1.91 |
| Clash score | 1.95 |
| C-beta outliers (%) | 0 |
| CaBLAM outliers (%) | 1.73 |
| CC (mask) | 0.82 |
| MolProbity score | 1.52 |
| EMRinger score | 3.75 |

